# Magnetically induced magnetosome chain (MAGiC): A biogenic magnetic-particle-imaging tracer with high performance and navigability

**DOI:** 10.1101/2024.12.04.626925

**Authors:** Lei Li, Chan Zhao, Zhiyuan Zhao, Dou Han, Yidong Liao, Yanjun Liu, Sijia Liu, Xiaohua Jia, Jiesheng Tian, Qing Liu, Xin Feng, Jie Tian

## Abstract

Magnetic particle imaging (MPI) enables real-time, sensitive and quantitative visualization of magnetic tracers’ spatial distribution, augmenting the capability of in vivo imaging technologies. Previous tracer studies in MPI have primarily focused on superparamagnetic nanoparticles; however, their non-ideal sigmoidal magnetization response limits the spatial resolution. Here we demonstrate the utilization of magnetically induced magnetosome chain (MAGiC) as a novel superferromagnetic MPI tracer, exhibiting a 25-fold improvement in resolution and a 91-fold enhancement in signal intensity compared to the commercial tracer VivoTrax+™. The spatial resolution of MPI was pushed to an unprecedented 80 μm under a 4 T/m gradient field. Additionally, MAGiC can be precisely controlled using magnetic fields, enabling it to function as a MPI trackable microrobot. We provided a theoretical model elucidating MAGiC’s unique properties, and validated its imaging and actuation performance through phantom studies and in vivo experiments. As a high-performance MPI tracer and magnetic microrobot with exceptional capabilities, MAGiC holds tremendous potential for diverse applications including cell tracking, targeted drug delivery as well as therapeutic interventions.

## Introduction

In translational research, *in vivo* imaging has become an essential tool, effectively bridging the gap between exploratory in vitro studies and clinical investigations^1–3^. Consequently, there is an increasing demand for advanced *in vivo* imaging techniques that offer highly sensitive, dynamic, and quantitative observations. Optical imaging, magnetic resonance imaging (MRI), and positron emission tomography (PET) are among the most commonly employed modalities for *in vivo* visualization. However, the effective and accurate visualization of objects using optical imaging is limited to a depth of 2 cm, posing challenges for achieving comprehensive and quantitative imaging beyond this range^4^. MRI is constrained by prolonged acquisition times and limited sensitivity when utilizing gadolinium or iron-based contrast agents^5^. PET, while exhibiting high sensitivity without depth limitations, is restricted by millimeter-scale resolution and its dependence on radioactive tracers^6^.

Magnetic Particle Imaging (MPI) is an emerging *in vivo* imaging modality that utilizes magnetic nanoparticles (MNPs) as tracers to directly visualize their spatial distribution within biological tissues^7^. MPI signals are generated from the nonlinear magnetization response of MNPs under alternating magnetic fields, which surpasses the proton magnetization signal in 7T MRI by a factor of 22 million^8^. Consequently, MPI offers unique advantages, including unrestricted depth penetration, high sensitivity, rapid imaging capabilities, radiation-free, and minimal interference from tissue background^9–11^. These characteristics have enabled MPI to demonstrate significant application potential in areas such as cell tracking^12,13^, cancer imaging^14,15^, cardiovascular imaging^16,17^, gastrointestinal imaging^18^, image-guided drug delivery^19^, and nanorobotics monitoring^20,21^.

Currently, the primary challenge encountered by MPI lies in its limited spatial resolution, which is compromised by the non-ideal sigmoidal magnetization response with commonly used superparamagnetic tracers^22,23^. Considerable research efforts have been directed towards enhancing the saturation magnetization of MPI tracers, which does not contribute to the improvement in resolution^8,12,24^. Recently, a pilot study introduced superferromagnetic nanoparticle chains, which significantly enhances the spatial resolution of MPI by an order of magnitude^23^ due to their square-shape magnetization response. However, the ultrahigh MPI performance of chemically synthesized nanoparticle chains is currently limited to organic solvents, restricting their feasibility for *in vivo* applications^23,25^. Therefore, the development of biocompatible superferromagnetic tracers is crucial for enhancing the performance of MPI, particularly in terms of resolution.

Magnetosomes, which are naturally biosynthesized MNPs composed of magnetite, are found in magnetotactic bacteria^26^. Previous studies have demonstrated their potential for MPI imaging, showing performance comparable to that of conventional MPI tracers^27,28^. Notably, magnetosomes exhibit square-shaped static hysteresis curves in vibrating sample magnetometry (VSM)^29^ when arranged in chains, resembling the superferromagnetic response reported in previous studies^23^. This observation motivated us to investigate the dynamic magnetization response of magnetosome chains and their potential for enhancing MPI performance.

In this study, we report that ‘magnetically induced magnetosome chain’ (MAGiC, Fig. 1a) exhibits superferromagnetic magnetization response under sinusoidal excitation, yielding a remarkable 25-fold improvement in resolution and a 91-fold enhancement in signal compared to the commercially available MPI tracer VivoTrax+^TM^ (Fig. 1b). This represents the highest resolution and one of the most sensitive MPI tracers reported to date. The MPI performance of MAGiC was validated through phantom studies and *in vivo* gastrointestinal imaging experiment. Furthermore, we demonstrated that MAGiC can be magnetically controlled, highlighting its potential as a MPI trackable microrobot (Fig. 1c, 1d). Overall, MAGiC is biosynthesized, exhibits exceptional MPI performance, possesses magnetic actuation capabilities, and demonstrates good biocompatibility. These distinctive characteristics establish it as a multi-functional nanoplatform with broad applications including cell tracking, imaging-guided drug delivery and therapeutic interventions.

**Fig.1.**
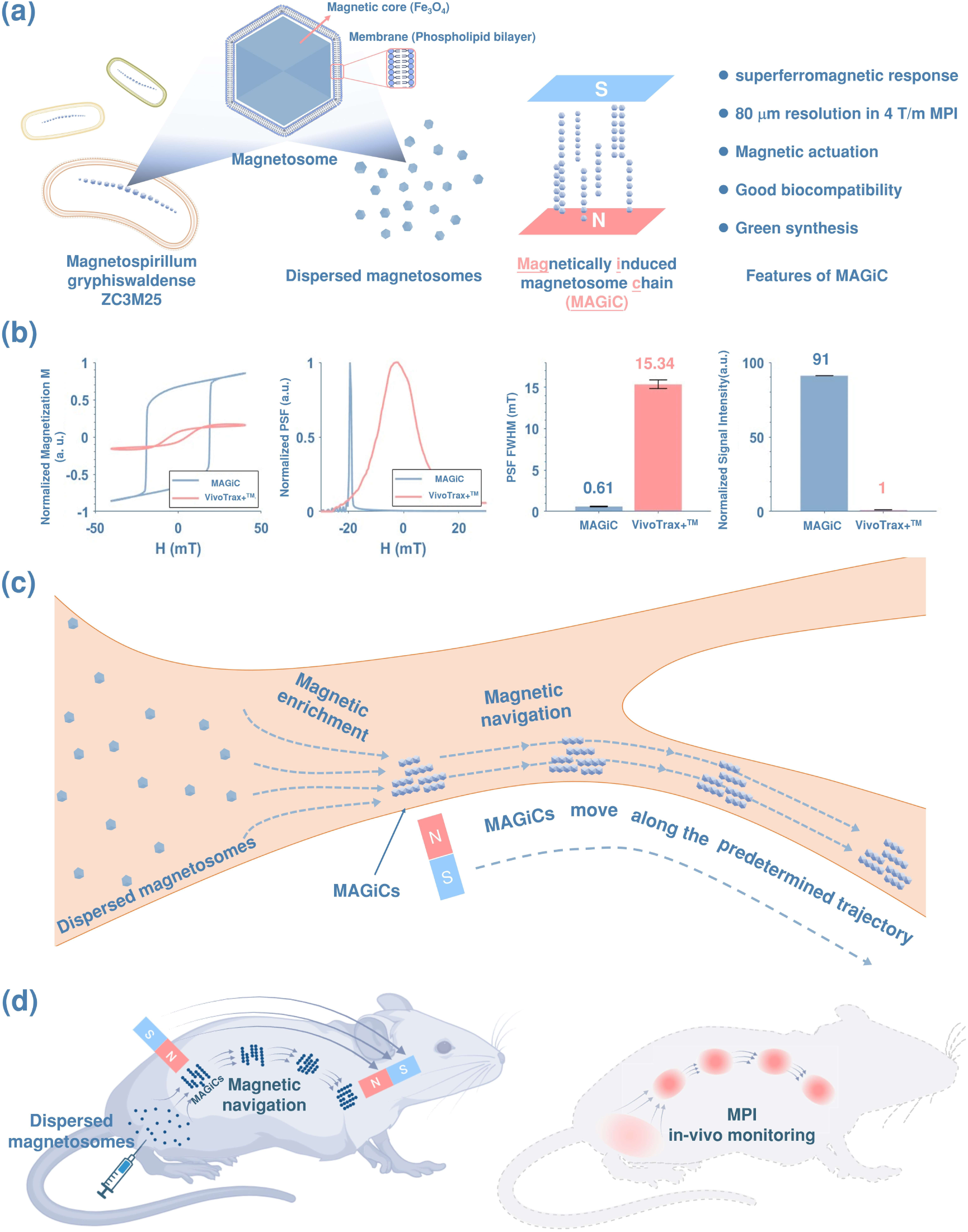
Schematics of the formation process, MPI performance, and magnetic actuation of MAGiC. (a) Illustration of magnetosomes purified from ZC3M25 assembling into MAGiCs under the induction of an external magnetic field. (b) The dynamic magnetization curve, normalized PSF, PSF FWHM, and normalized signal intensity of MAGiC and VivoTrax+^TM^. MAGiC exhibits exceptional MPI performance due to its superferromagnetic response under alternating magnetic fields, resulting in a 25-fold improvement in resolution and a 91-fold enhancement in signal intensity compared to superparamagnetic tracer VivoTrax+^TM^.(n = 10) (c) Illustration of the magnetic enrichment and navigation of MAGiC. (d) Illustration of MPI-guided navigation of MAGiC in vivo.

## Results

### Synthesis and characterization of MAGiC

The magnetotactic bacterium Magnetospirillum gryphiswaldense ZC3M25 (Fig. 2a) was cultivated and employed for the biosynthesis of magnetosomes^30^. Isolated magnetosomes were harvested from the bacterial cells using sonication, which consist of cuboctahedron-shaped magnetite core with average diameter approximately 27 nm (Fig. 2b). Surrounding the magnetite core was a membrane with a thickness of 3.6 nm. The zeta potential measurement yielded -31.3 ± 5.67 mV. VSM measurements demonstrated that the magnetosomes had a saturation magnetization of approximately 39 emu/g (Fig. 2c). The isolated magnetosomes exhibited good dispersion properties and maintained good colloidal stability in solution (Fig. 2d and 2e). The dynamic magnetization response of the dispersed magnetosomes (Fig. 2f) resembled that of single-core MNPs^31^. The full width at half maximum (FWHM) of their PSF was measured to be 13.5 mT (Fig 2g). Intriguingly, following a uniform magnetic field induction (80 mT), the magnetosomes assembled into MAGiCs, as depicted in the TEM image (Fig. 2h). The substantial aggregation seen in Fig. 2h does not represent their native state in solution, as the drying process exacerbated aggregation during sample preparation.

**Fig. 2.**
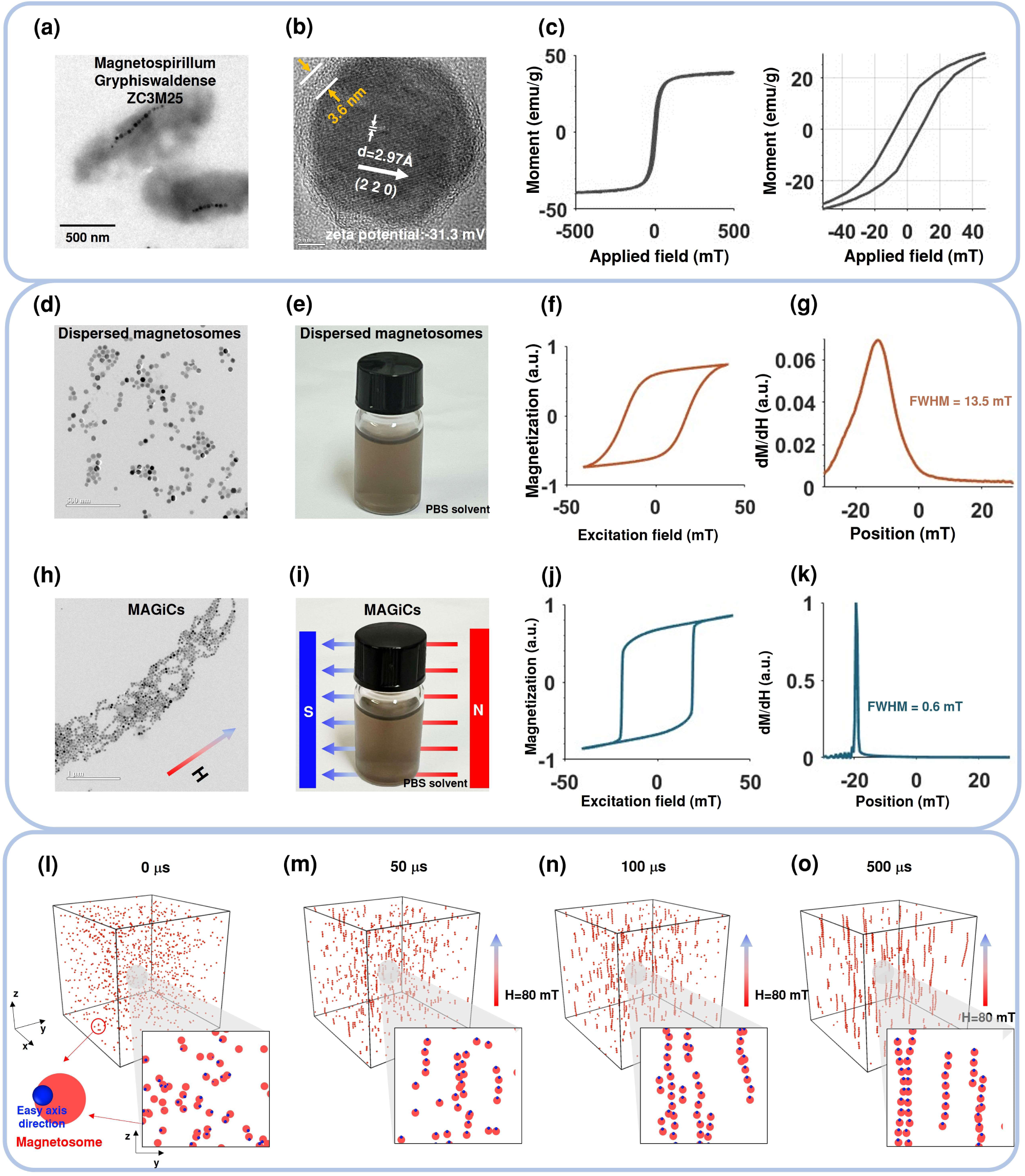
Characterization of dispersed magnetosomes and MAGiC. (a) TEM image of Magnetospirillum gryphiswaldense ZC3M25 with magnetosomes located inside the cellular body. (b) TEM image of a purified magnetosome. (c) Static magnetization curve of magnetosomes recorded using VSM. (d) TEM image of the dispersed magnetosomes. (e) Photograph depicting dispersed magnetosomes in PBS. (f) The dynamic magnetization curve of dispersed magnetosomes measured with MPS. (g) MPI PSF of dispersed magnetosomes. (h) TEM image of MAGiCs orientating towards the magnetic field. (i) Photograph of MAGiCs in PBS. (j) The dynamic magnetization curve of MAGiC measured with MPS, revealing superferromagnetic response. (k) MPI PSF of MAGiC. (l)-(o) Simulated MAGiC formation process under the magnetic field induction.

Hydrodynamic size of MAGiC was 558 ± 114 nm, indicating about 10-15 magnetosomes in each chain (Fig. S1). As demonstrated in Fig. 2i, MAGiC remained good colloidal stability in solution after brief magnetic induction. However, unlike the isolated magnetosomes (Fig. 2f), MAGiC exhibited superferromagnetism, characterized by a square-shaped dynamic hysteresis loop (Fig. 2j) and a considerably narrower PSF (Fig. 2k), indicating a substantial enhancement in both MPI signal and resolution. The formation process of MAGiC was simulated and visually demonstrated in Mov. 1 and Fig. 2l-o. Prior to magnetic induction, dispersed magnetosomes displayed random orientations of their easy axes (Fig. 2l). Upon application of a magnetic field, the easy axes of the magnetosomes aligned with the field direction, leading to enhanced magnetic interactions between adjacent magnetosomes (Fig. 2m) and subsequent formation of MAGiCs (Fig. 2n, 2o) that aligned parallel to the applied field^32^. In addition to simulations, we conducted experiments with different induction times and demonstrated that 5 s of induction at 80 mT was sufficient for superferromagnetic MAGiC formation (Fig. S2).

### Physical model of MAGiC’s superferromagnetic response

Within magnetotactic bacteria, cytoskeleton proteins organize magnetosomes into chains; however, these chains do not exhibit superferromagnetic responses like MAGiC despite their morphological similarity (Fig. S3). This observation prompted us to explore the physics underlying the phenomenon of superferromagnetism in MAGiC. Unlike magnetosomes in bacterial cells that are attached to cytoskeletal filaments, free-floating magnetosomes in MAGiC can rotate and align with external magnetic fields through Brownian relaxation. Therefore, we propose that MAGiC’s superferromagnetism response can be attributed to a collective Brownian relaxation of individual magnetosomes. Indeed, hindering Brownian relaxation by increasing solvent viscosity attenuated the superferromagnetic behavior of MAGiC (Fig. S4).

The dynamic magnetization response of MAGiC is further elucidated in Fig. 3a, where a magnetization curve is presented alongside the depiction of particle orientations within MAGiC. Initially, as the excitation magnetic field increased from zero to its maximum (point A to B), the easy axes of individual magnetosomes aligned with the field direction, resulting in a fully ordered chain at point B. Subsequently, as the excitation field decreased to zero and reversed direction (from point B to C), interparticle interactions impeded the response of magnetosomes within MAGiC. Once the coercive force was surpassed in the reverse direction, the excitation field triggered a cooperative Brownian relaxation of magnetosomes within MAGiC, resembling a cascading effect originating from the chain ends and propagating toward the center^29^. This process resulted in complete alignment and reverse magnetization at point F. Fig. 3b shows the excitation field and received signal during this response process. The duration between points C and F was extremely short; however, all magnetosomes in MAGiC underwent collective Brownian relaxations, producing a sharp response signal during this period. This superferromagnetic magnetization response leads to an order of magnitude enhancement in MPI performance for MAGiC (Table S1), as measured by magnetic particle spectroscopy (MPS).

**Fig. 3.**
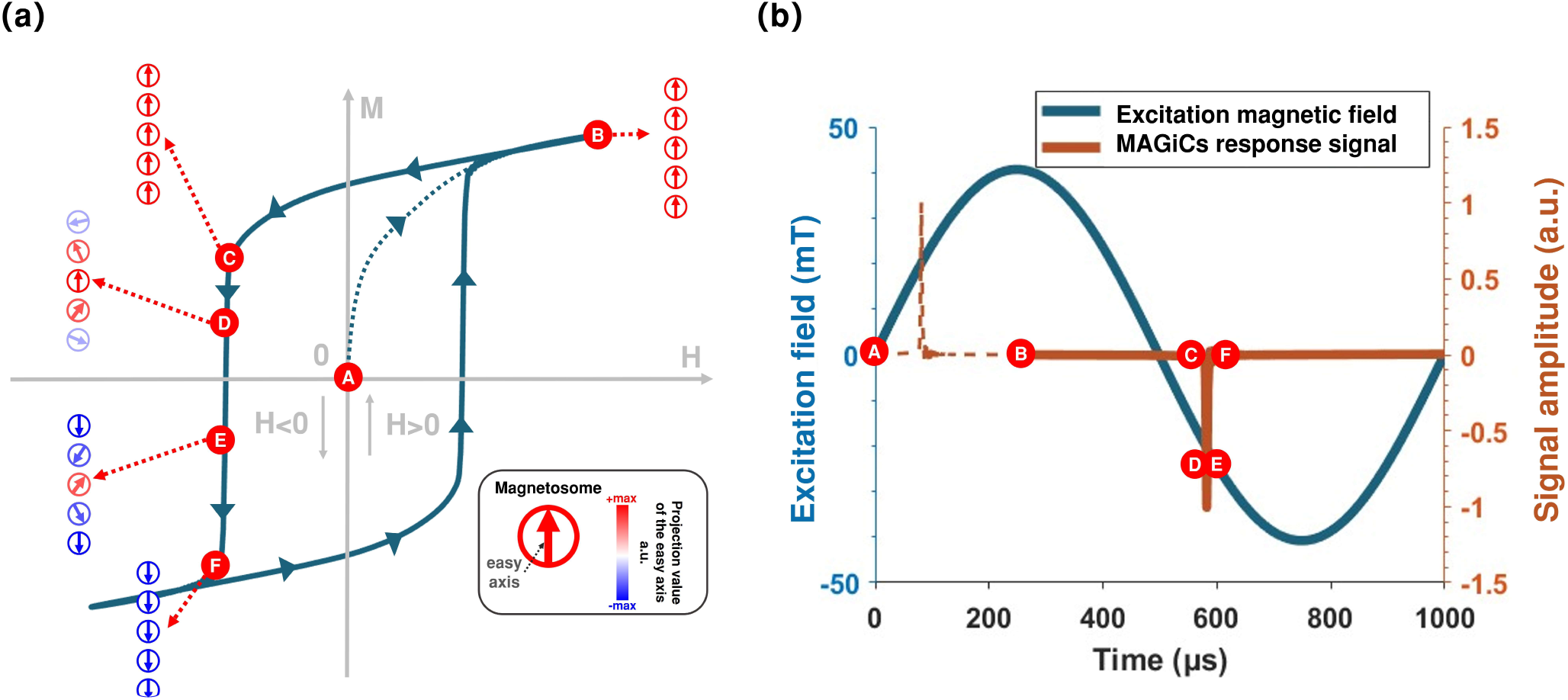
Superferromagnetic response of MAGiC under a sinusoidal excitation field. (a) M-H curve of MAGiC. Circles with arrows represent the magnetosomes, and their colors indicate the projection value of the magnetosome’s easy axis in relation to the magnetic field direction. Positive projections are depicted in red, while negative ones are shown in blue. (b) The excitation magnetic field waveform and the MAGiC response signal. The process between points C and F exhibits a very short duration but elicits a robust collective response, which significantly enhances both resolution and signal intensity in MPI. The signal curve between A and B was marked with dashed line, since it represents the steady state signal and does not correspond to the response during the A-B process in (a).

According to our physical model, the superferromagnetism of MAGiC arises from the collective Brownian relaxation of individual magnetosomes in response to external fields and their interparticle interactions. Therefore, we hypothesized that the superferromagnetism of MAGiC would be sensitive to factors such as magnetosome sizes^33^, interparticle interactions (Fig. 4a), and excitation field parameters. Computational simulations and experimental investigations were conducted to explore their influence and validate our hypothesis. As discussed in the supplementary file, an algorithm that couples the Landau-Lifshitz-Gilbert (LLG) equation into Brownian dynamics simulations was developed, accounting for both rotational and translational motion of the magnetosomes in MAGiC.

**Fig. 4.**
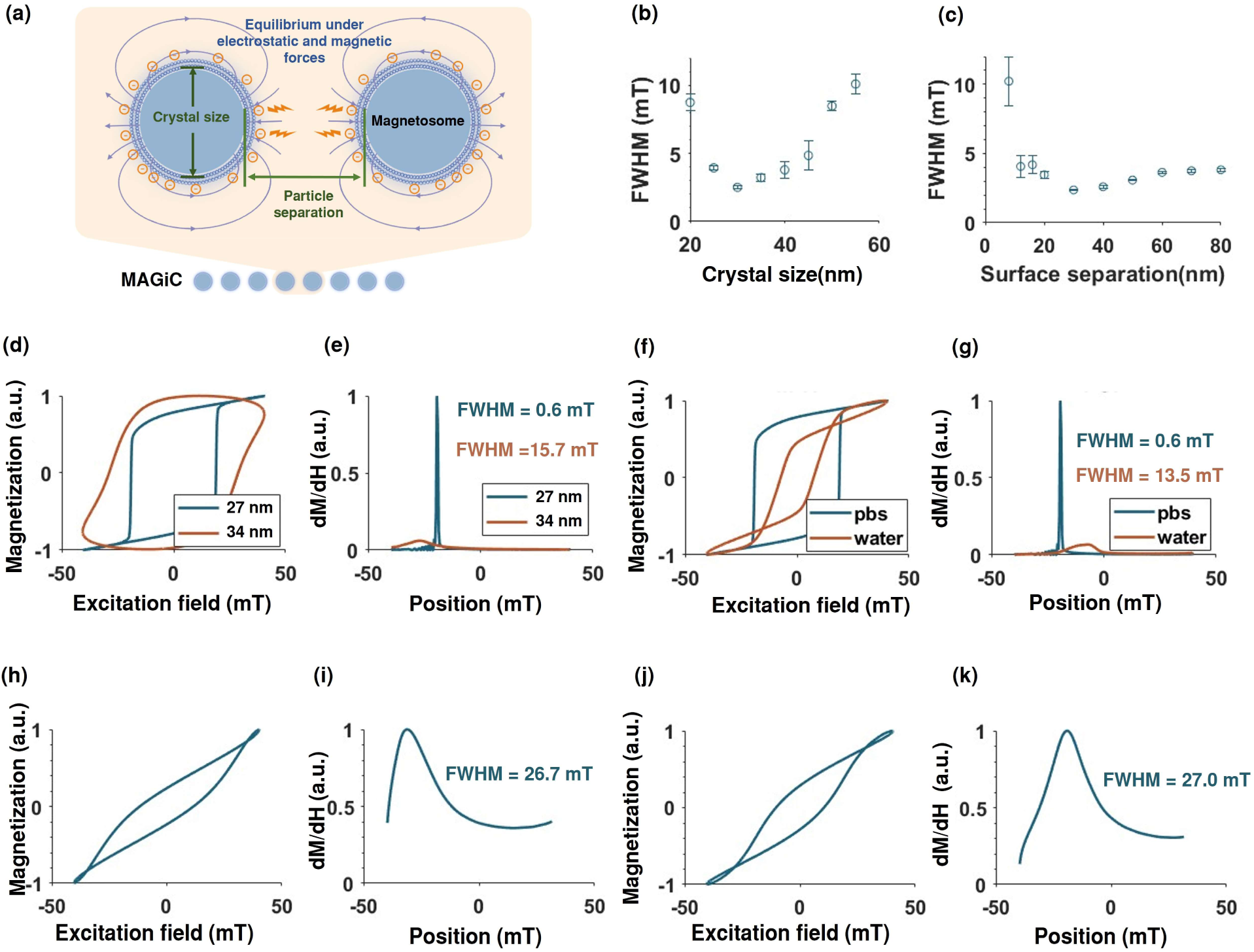
Factors affecting the MPI performance of MAGiC. (a) Schematic diagram of the interactions between neighboring magnetosomes in MAGiC; magnetic and electrostatic forces maintain a specific separation between them. (b) Dependence of PSF FWHM on the crystal size of magnetosomes based on simulation (n= 5). (c) Dependence of PSF FWHM on the surface separation between magnetosomes based on simulation (n= 5). (d) Dynamic magnetization responses of MAGiCs composed of 27 nm and 34 nm magnetosomes measured by MPS. (e) The normalized PSF of MAGiCs composed of 27 nm and 34 nm magnetosomes. (f) Dynamic magnetization response of MAGiCs in deionized water and PBS measured by MPS. (g) The normalized PSF of MAGiCs in deionized water and PBS. (h) Dynamic magnetization response and PSF (i) of MAGiC after the removal of the magnetosome membrane. (j) Dynamic magnetization response and PSF (k) of MAGiC after the removal of the magnetosome membrane proteins.

### Dependence of MAGiC’s MPI performance on crystal sizes

Based on the simulation results, MAGiC’s FWHM (inversely related to MPI performance) exhibits a non-monotonic trend as the crystal diameter increases from 20 nm to 55 nm (Fig. 4b), reaching an optimum around 30 nm. For crystal sizes below 25 nm, the diminished dipolar interaction cause MAGiC to transition into a superparamagnetic state; whereas for crystal sizes exceeding 35 nm, the increased Brownian relaxation time hinders rapid response of MAGiC to external magnetic fields, thereby compromising their MPI performance in both scenarios.

In the experiments, magnetosomes with optimal (27 nm) and suboptimal (34 nm) crystal diameters were obtained using ZC3M25 under different bacterial culture conditions (Fig. S5). Consistent with the simulation result, MAGiCs with different crystal sizes exhibited distinct magnetization behaviors (Fig. 4d and 4e). MAGiC composed of 34 nm magnetosomes displayed a significantly broadened PSF and inferior MPI performance compared to those composed of 27 nm magnetosomes.

### Dependence of MAGiC’s MPI performance on interparticle interactions

Neighboring magnetosomes in MAGiC exhibit a specific separation owing to a delicate balance between electrostatic repulsion and magnetic attraction (Fig. 4a). Our simulation showed that FWHM of MAGiC initially decreased and then increased as particle separations increase, with an optimal separation observed at approximately 30 nm (Fig. 4c). Beyond this optimum, as interparticle interaction diminishes, the MAGiC transits from superferromagnetic to superparamagnetic. Conversely, the strong interparticle interactions constrain the Brownian relaxation when magnetosomes are in close proximity, thereby impeding MAGiC’s prompt response to external fields.

To validate the simulation results, the interparticle interaction was adjusted by manipulating the solvent composition in the experiment. As previously reported, the presence of electrolytes in PBS enhances interparticle interactions compared to deionized water^34^. Indeed, transitioning from PBS to deionized water as the solvent resulted in a broadened PSF and deteriorated MPI performance, due to diminished interparticle interactions (Fig. 4f and 4g). Furthermore, removal of magnetosome membrane and its associated proteins was performed to further increase interparticle interactions, which again led to a substantial reduction in MPI performance (Fig. 4. h-k), consistent with our simulation. Based on these observations, membrane-intact MAGiCs with a crystal size of 27 nm were dispersed in PBS for subsequent imaging experiments.

### Effect of the excitation magnetic field parameters on MAGiC’s MPI performance

In addition to the particle size and interparticle interaction, the MPI performance of MAGiC also varies with excitation field parameters. In our simulation (Fig. 5a), enhanced MPI performance was observed only when the excitation field amplitudes exceeded the coercivity threshold (10 mT in Fig. 5a). Below this threshold, MAGiC cannot be fully magnetized, resulting in a weak and delayed magnetic response. Additionally, MPI performance declines with increased excitation field frequency (Fig. 5b) due to the relaxation effect^22,33^.

**Fig. 5.**
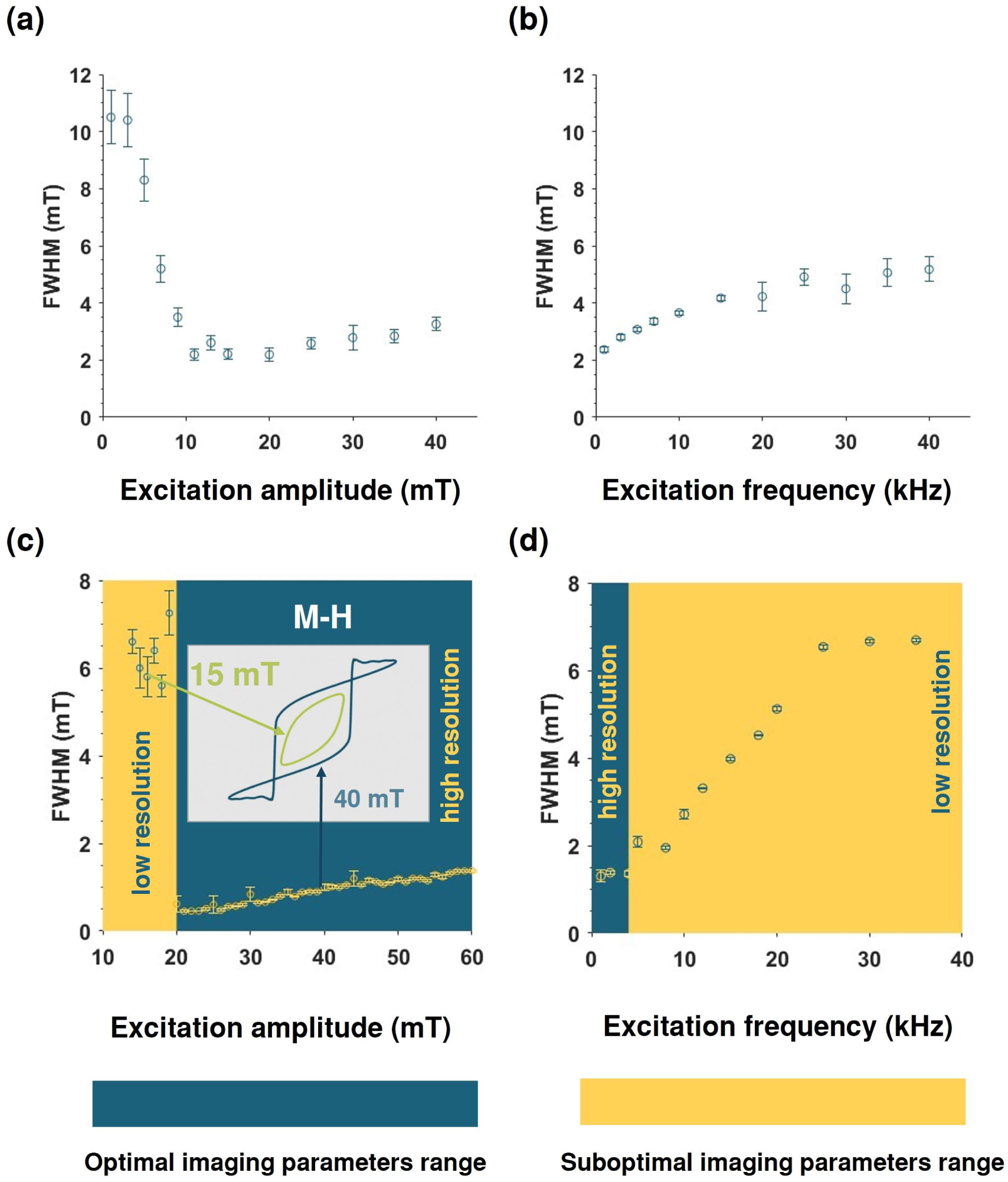
MPI performance of MAGiC under varying excitation field parameters. Dependence of FWHM on (a) excitation field amplitudes and (b) frequencies based on simulation (n=5). Dependence of FWHM on (c) excitation field amplitudes and (d) frequencies measured using MPS (n=10). Optimal and suboptimal parameter ranges are denoted by blue and yellow markings, respectively.

MPS measurements demonstrated a consistent trend with slightly different threshold values, which may be attributed to the discrepancy in simulation parameters. In Fig. 5c and 5d, the measured optimal excitation parameters were highlighted with a blue background, while the suboptimal parameter range was indicated with a yellow background. Based on these findings, excitation parameters of 1 kHz and 40 mT were selected for subsequent imaging experiments. Given that existing commercial MPI scanners utilize high-frequency (25 kHz-45 kHz) excitation fields unsuitable for MAGiC, dedicated MPI scanners were developed with optimized excitation parameters tailored specifically for MAGiC imaging (Fig. S6, S7).

### Phantom Imaging Results

The phantom imaging results of MAGiC and the commercial tracer synomag®-D are compared in Fig. 6. To ensure a fair comparison, synomag®-D images were acquired using a commercial MPI scanner (Momentum, Magnetic Insight Inc.). As shown in Fig. 6b, the MAGiC image successfully distinguishes point sources with 0.3 mm spacing in the excitation direction (x-direction), whereas the synomag®-D images fail to resolve points with spacing below 1 mm. Notably, the field gradient in the Momentum scanner (5.7 T/m) was 4.5 times higher than that of the MAGiC scanner (1.25 T/m). Accounting for these field gradients, MAGiC demonstrated a remarkable 15-fold improvement in resolution compared to synomag®-D. The “MPI” letter phantom images on the right side of Fig. 6 clearly demonstrate that MAGiC produces sharper images with more details than those obtained using synomag®-D. The 3D imaging results are shown in Fig. S8. To further evaluate the imaging performance of MAGiC at higher field gradients, its resolution was examined using line pair phantoms on a dedicated 4 T/m 1D scanner (Fig. S9, S10). An impressive resolution of 80 μm was achieved, surpassing those of synomag®-D and VivoTrax+^TM^ by an order of magnitude (Fig. S9, S10). To the best of our knowledge, this value represents the highest resolution in MPI reported to date.

**Fig. 6.**
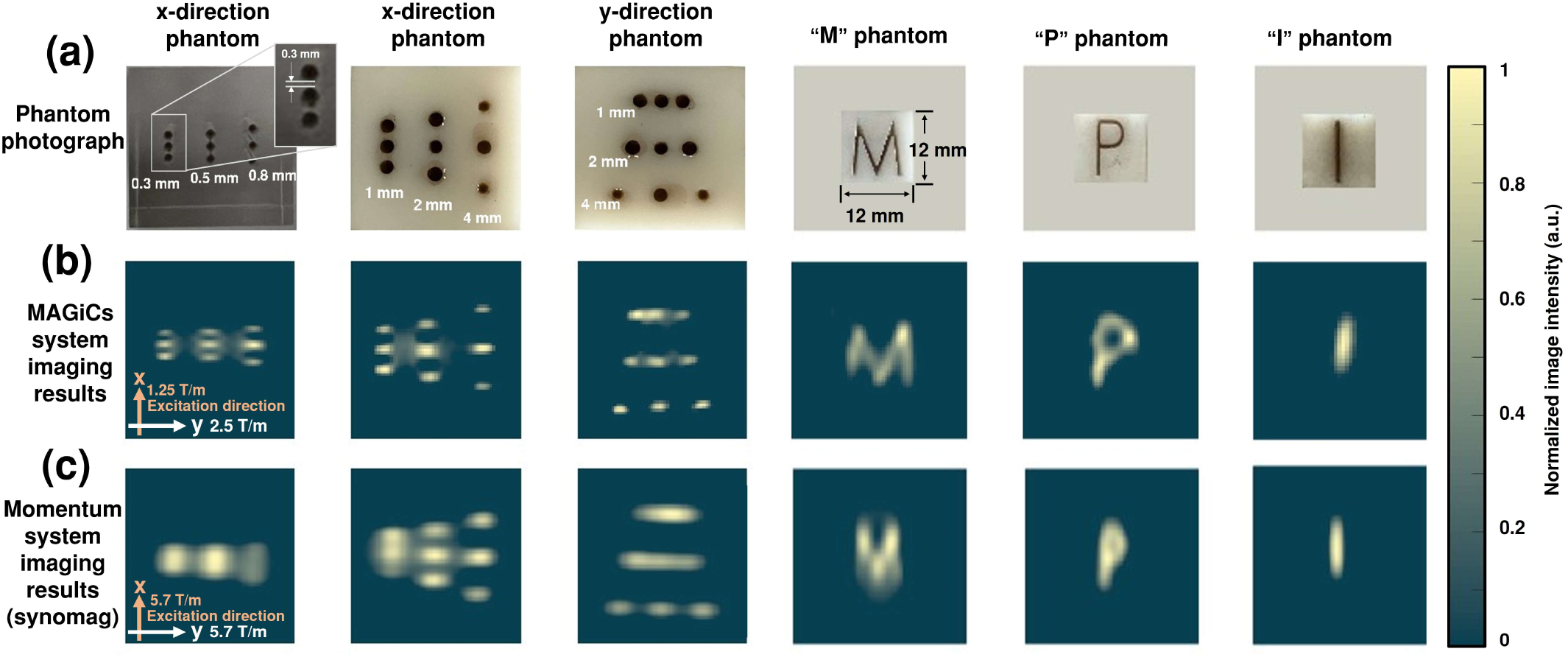
Phantom imaging results of MAGiC and synomag®-D. (a) The photograph of resolution phantoms and letter phantoms used in the study. (b) Imaging results of MAGiC acquired with the in-house built MPI scanners dedicated for MAGiC, revealing a maximum resolution of 0.3 mm under a gradient of 1.25 T/m in the x-direction. The spatial resolution in the y-direction is 1 mm. c) Imaging results of synomag®-D acquired with the Momentum MPI scanner, demonstrating a maximum resolution of 1 mm under a gradient of 5.7 T/m.

### *In Vivo* Imaging of MAGiC in gastrointestinal tract

Recently, engineered nanoplatforms have been utilized for active drug delivery and localized therapy in the gastrointestinal (GI) tract^35,36^. However, real-time tracking of nanoplatforms within the GI tract *in vivo* remains challenging owing to limited imaging depth and prolonged acquisition time associated with existing imaging techniques. We proposed that MAGiC could serve as a novel multifunctional nanoplatform with both delivery and imaging capabilities, enabling quantitative real-time tracking using MPI. As a proof of concept, SD rats were orally administered encapsulated MAGiCs, and MPI images were acquired at multiple time points to monitor the release, retention and clearance dynamics of MAGiCs within the GI tract (Fig. 7a). The enteric capsule^35^ containing MAGiCs was imaged in vitro prior to administration, and the amount of MAGiCs was quantified based on the signal intensity (Fig. 7b). As shown in Fig. 7c, the enteric capsule remained intact in the stomach during the initial 2 h post-administration. At 3 h, the capsule traversed into the intestine, where it gradually dissolved and released MAGiCs. A decrease in image signal intensity accompanied by an expansion of the signal area was observed. From 3 h to 24 h post-gavage, MAGiCs progressively distributed and moved throughout the GI tract, with a continuous decline in image signal intensity. After 24 h, most of the administered MAGiCs were cleared, while approximately 24% remained within the lower GI tract. These findings demonstrate the feasibility and efficacy of utilizing MPI imaging for dynamically tracking of MAGiCs within deep tissues *in vivo*. Meanwhile, blood tests and histopathological examinations were conducted at both 24 h and 7 days post MAGiCs administration. There were no significant variations observed in terms of liver and kidney functions (Fig. S11), as well as the morphologies of major organs or intestinal tissues when compared to those of the control group (Fig. S11-S12). These findings confirm the good biocompatibility associated with the utilization of MAGiC nanoplatforms.

**Fig. 7.**
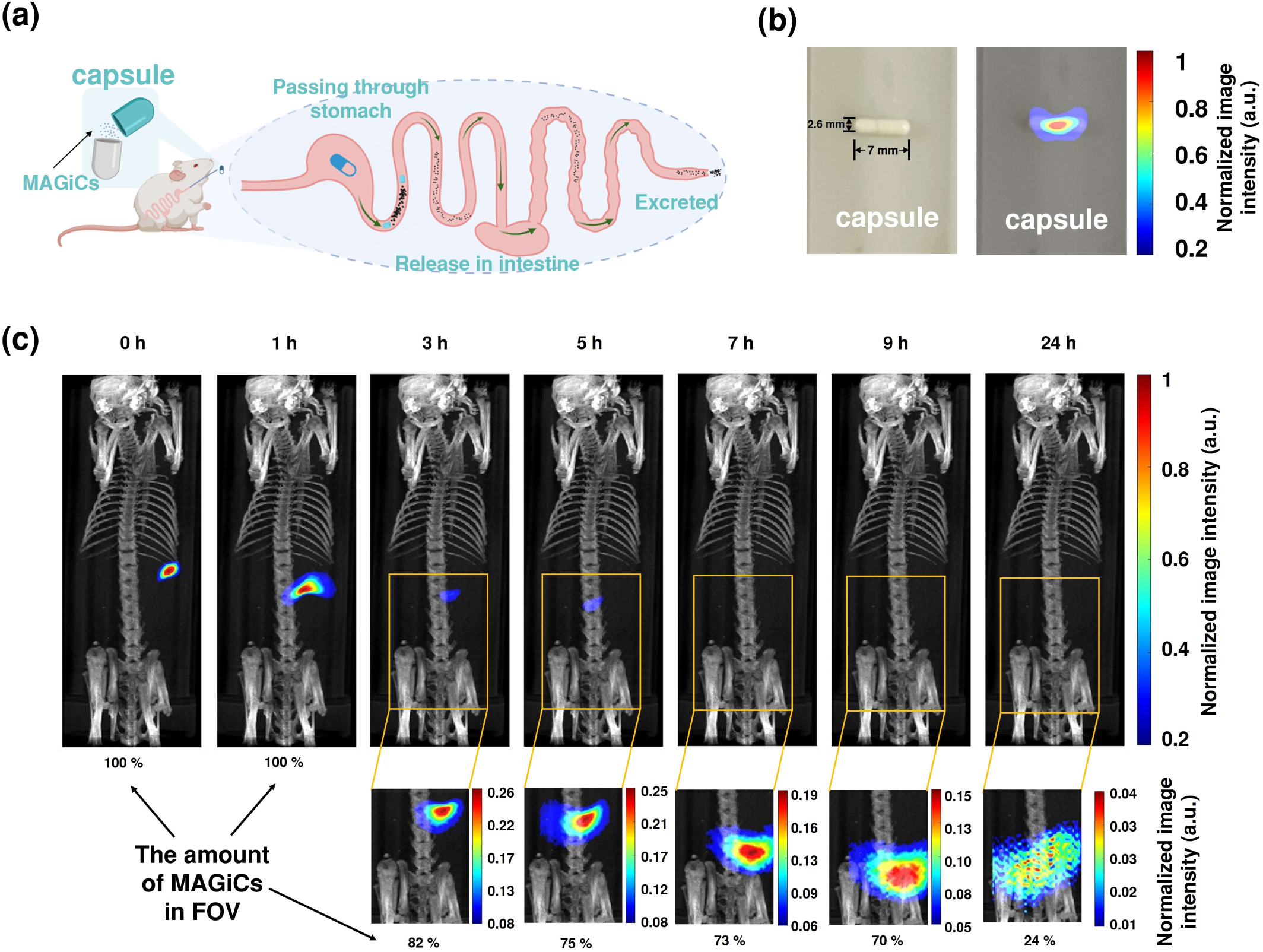
In vivo MPI tracking of the MAGiC capsule administered via oral gavage. (a) Illustration demonstrating the journey of the MAGiC capsule through the gastrointestinal tract after oral administration. The capsule traverses the stomach, dissolves in the intestine, and eventually gets excreted. (b) The photograph and MPI image of the enteric-coated capsule with a hydrophobic inner layer, filled with 10 μL of 5 mg/mL MAGiC. (c) In vivo MPI tracking of the MAGiC capsule after administration. The upper row displays the imaging results from 0 to 24 hours using a consistent colorbar, while images with optimized colorbars are shown in the lower row for enhanced visualization. (n= 3 rats).

### Magnetic actuation of MAGiC

In addition to its exceptional MPI performance, we demonstrated the capability of MAGiC to be actuated by magnetic fields, enabling it to function as a microscale magnetic robot. As a proof of concept, we first showcased the targeted enrichment of MAGiC in solution using a permanent magnet, achieving an impressive enrichment rate of approximately 94% within just 50 s (Fig. S13). Subsequently, we investigated the locomotion of a cluster of MAGiC along predetermined paths guided by a permanent magnet (Fig. S14, Mov.2). Furthermore, we successfully demonstrated the upstream navigation capability of MAGiC in fluidic environments with flow rates of up to 20 mL/min (Fig. S14, Mov.3).

The *in vivo* magnetic actuation of MAGiC was also investigated. As illustrated in Fig. 8a, MAGiCs were administered intraperitoneally to SD rats and guided to different locations using a magnet. Simultaneously, the distribution and locomotion of MAGiCs were quantitatively tracked through MPI. The images show that within 5 min after administration, MAGiCs rapidly dispersed throughout the abdominal cavity (Fig. 8b, time point 1). Subsequently, a magnet was positioned at target site 1 to facilitate the recruitment of MAGiCs. After 80 min, approximately 76% of the injected MAGiCs were recruited to target site 1 (Fig. 8b, time point 2). The magnet was then repositioned to enable the gradual migration of MAGiCs towards the second target location. After 80 min, approximately 28% of the MAGiCs reached target site 2, and their migration route was clearly identified in the image (Fig. 8b, time point 3). Finally, after 220 min, nearly all the recruited MAGiCs successively reached site 2 (Fig. 8b, time point 4). The biosafety of intraperitoneal administration of MAGiC was assessed through blood tests and histopathological examinations (Fig. S15-S16). Comparisons with the control group revealed no notable changes in liver and kidney functions nor in the morphologies of major organs.

**Fig. 8.**
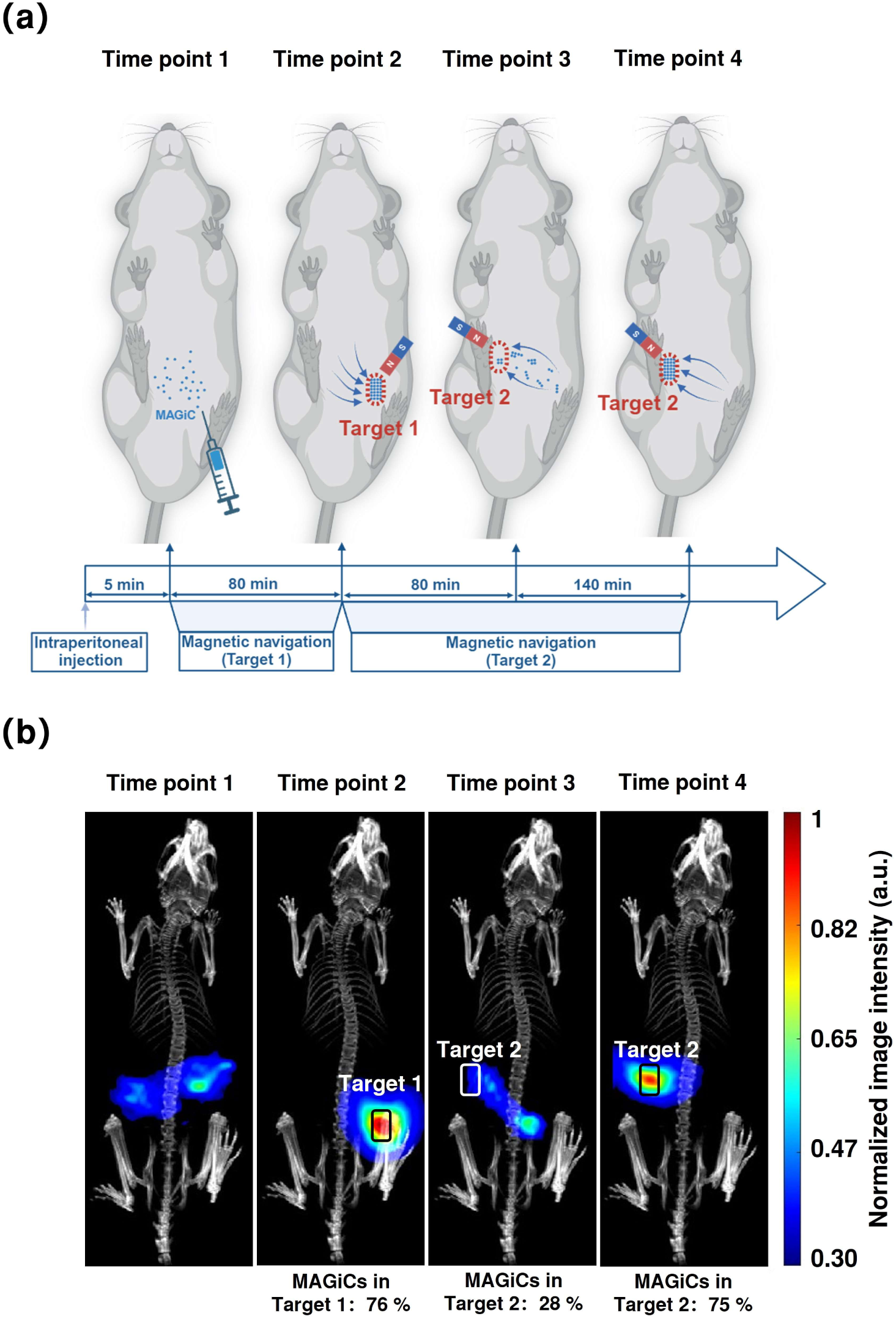
MPI guided magnetic actuation of MAGiC in vivo. (a) Schematic diagram of the experimental procedure. (b) In vivo MPI monitoring of the navigation process. Upon administration, the MAGiCs were dispersed within the abdominal cavity. After an enrichment period of 80 minutes, approximately 76% of MAGiCs reached the vicinity of target position 1. During navigation toward position 2, approximately 28% and 75% of MAGiCs successfully reached the target position at intervals of 80 minutes and 220 minutes, respectively (n=2 rats).

## Discussion

MPI exhibits unrestricted tissue penetration, exceptional sensitivity, and non-radiative properties, establishing it as one of the most promising *in vivo* imaging modalities for clinical translation. However, the resolution of MPI consistently poses a bottleneck challenge in the field. In this work, we introduce MAGiC, a novel biogenic MPI tracer that exhibits exceptional sensitivity with a detection limit of 6.8 ng (Fig. S17) and achieves an impressive spatial resolution of 80 μm at 4T/m - representing the highest reported resolution in the field of MPI to date. Additionally, a physical model was developed to elucidate the distinctive superferromagnetic response of MAGiC, while identifying key factors influencing its exceptional property. These findings establish a fundamental basis for future applications and advancements of MAGiC.

Comparing to previously reported chemically synthesized superferromagnetic particle chains^23^, MAGiC demonstrates a threefold improvement in resolution. Importantly, the superferromagnetism of MAGiC remains stable in PBS, whereas previously reported particle chains only exhibit superferromagnetic behavior in organic solvents^23,25^. Therefore, MAGiC effectively addresses challenges related to toxicity when used in vivo.

Moreover, what distinguishes MAGiC is its biogenic nature. Unlike hydrothermal synthesis methods for MNPs, which require high temperatures and pressures, biosynthesis of magnetosomes occurs under mild conditions within bacterial cells on a comparable time scale^37^. Under stringent gene regulation, bacteria consistently synthesize magnetosomes with uniform crystal structures, compositions, and sizes^26,38^. Additionally, the utilization of fermenter culture enables mass production of magnetosomes with consistent quality^39,40^, effectively resolving the scalability problem frequently encountered in chemical synthesis. Our findings also demonstrate good biocompatibility of magnetosomes, aligns with previous reports^41,42^.

In addition to its exceptional MPI performance, MAGiC can be effectively actuated by external magnetic fields, positioning it as a highly promising candidate for the development of trackable magnetic microrobots. Currently, the limited application of microrobot in vivo is attribute to the lack of real-time tracking techniques. We demonstrate the *in vivo* navigability and enrichment of MAGiC, with accurate tracking and quantification using MPI. These findings highlight the potential of MAGiC for applications such as image-guided active drug delivery and therapy.

In this study, we compared the performance of two types of MAGiC with average crystal sizes of 27 nm and 34 nm, with the smaller one exhibits significantly enhanced MPI performance. Given the genetic regulation of magnetosome synthesis, future work could leverage gene editing techniques to produce magnetosomes with diverse crystal shapes and sizes for better performance. Furthermore, it may be feasible to directly synthesize dispersed magnetosomes by deleting the mamJ or mamY gene^28^, thereby eliminating the need for subsequent purification steps. Besides, Previous studies have reported successful surface modifications of magnetosomes^43^. Future investigations may enable the functionalization of MAGiC through molecular targeting or therapeutic agents, thereby establishing it as a versatile nanoplatform for disease diagnosis and targeted therapy.

In summary, the biogenic MAGiC exhibits superferromagnetism and exceptional MPI performance, outperforming current tracers in terms of sensitivity and resolution, while effectively addressing the issues of biocompatibility and scalability. Additionally, MAGiC can be actuated by a magnetic field, highlighting their potential as magnetic microrobots. Our findings validate the real-time and quantitative imaging capabilities of MAGiC in deep tissue, as well as the MPI-guided magnetic actuation in vivo. Thus, MAGiC represents a multifunctional nanoplatform capable of facilitating high performance MPI imaging, efficient magnetic actuation, drug loading, and hyperthermia applications. These capabilities demonstrate significant potential for disease diagnosis, image-guided active drug delivery and therapy.

## Data availability

The main data supporting the findings of this study are available within the paper and its supplementary material. Source data are provided with this paper.

## Competing interests

The authors declare no competing interests.

## Author Contributions

Lei Li and Xin Feng were responsible for the construction hardware, the algorithms, data processing, and imaging experiments. Chan Zhao, Dou Han, Qing Liu and Jiesheng Tian contributed to the cultivation of magnetotactic bacteria and processing. Zhiyuan Zhao developed the simulation framework. Yidong Liao, Yanjun Liu, Sijia Liu and Xiaohua Jia were responsible for control system, phantoms and biocompatibility. Jie Tian supervised the project. All authors edited and approved the submitted version of the manuscript.

## Acknowledgements

This work was supported in part by the National Natural Science Foundation of China under Grant 62027901, Grant 32350010, Grant 82302407, Grant 81227901, and No. 12304247, in part by Beijing Natural Science Foundation under Grant 7232346 and Grant L234056, in part by Beijing Nova Program under Grant 20240484528. The authors would like to acknowledge the instrumental and technical support of Multimodal Biomedical Imaging Experimental Platform, Institute of Automation, Chinese Academy of Sciences.

## Methods and Materials

### Bacteria Culture

A mutant strain of magnetotactic bacteria, *Magnetospirillum gryphiswaldense* ZC3M25 (CGMCC 32901, China General Microbiological Culture Collection Center, Beijing, China) was used in this work. Bacteria were cultured in bottles or fermenters to synthesize magnetosomes of varying sizes. To synthesize magnetosomes with a crystal diameter of 27 nm, ZC3M25 was cultured in screw-capped bottles at 30 ℃ and 100 rpm using a sodium lactate medium. Magnetosome synthesis was induced by adding 60 μM ferric citrate (CAS#3522-50-7, Yuanye Biotech, Beijing, China) and shaking at 30 ℃ and 100 rpm for 24 h.

To synthesize magnetosomes with 34 nm crystal size, ZC3M25 was inoculated at 10% (v/v) into 500 mL sodium lactate medium as seed solution and transferred to a 7.5 L fermenter. Fermentation medium and feed medium were determined previously^30^. Fermentation culture was performed with initial air flow rate of 0.5 L/min at 30 ℃ and 100 rpm. Once the dissolved oxygen concentration (dO_2_) decreased to 15%, the airflow rate was increased to 1 L/min. The dO_2_ concentration was then maintained between 0-1% by regular agitation (adding 20 rpm) every 2 hours. pH=7.0 was maintained by automatically replenishing the feed medium. Cell growth (OD_565_) and magnetic response (Cmag) were measured at 4-hour intervals.

### Preparation of Magnetosomes

Bacteria was collected by centrifugation at 5000 rpm for 10 min. The collected bacteria cells were resuspended with 10 mM PBS in a beaker and were disrupted by an Ultrasonic cell crusher (SCIENTZ-IID, Ningbo Scientz Biotechnology Co., Ltd., Ningbo, China). Specifically, the following settings were used: 300 W power output, 3 s each time, 5 s working interval, and a total working time of 60 min. After ultrasonication, the beaker was placed on a magnet at 4 ℃ overnight. Supernatant was discarded, and the sediments were resuspended in PBS. The above-mentioned ultrasonication and magnetic collection processes were repeated 7-8 times, with ultrasonication power and time gradually decreasing. OD_260_ and OD_280_ of the supernatant were measured to determine the protein concentration C: C (mg/mL) = -1.45 × OD_260_-0.74 × OD_280_. Magnetosome purification process was considered as completed when the protein concentration of the supernatant is lower than 0.1 mg/mL.

### Characterization of magnetosomes and MAGiCs

The morphology of MAGiC and dispersed magnetosomes were examined using transmission electron microscopy (TEM, JEM 1200 EX, jeol, Tokyo Japan). Crystal sizes of magnetosomes were measured from the TEM images. The static magnetization curve was measured using a vibrating sample magnetometer (VSM-220, Yingpu Magnetoelectric, Changchun, China). The hydrodynamic size and zeta potential were examined using a dynamic light scattering Zetasizer (Nano-ZS90, Malvern Instruments, Malvern, UK).

### Simulation methods

In current work, we developed the algorithms that couple the Landau-Lifshitz-Gilbert (LLG) equation into Brownian dynamics simulations by considering both rotational and translational motion of the magnetosomes. The magnetosomes were modeled by uniform spherical particles consisting of homogeneous polymer shell and single-domain magnetic core with cubic magnetocrystalline anisotropy (which is typical for magnetite). The damped precession of the magnetic dipole moment towards the effective field **B**^(*i*)^_eff_ can be described by the LLG equation:

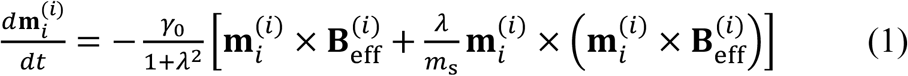

where the superscript (*i*) denotes the vectors in the coordinates of particle *i* (= 1, …, *N*), *N* represents the total number of the particles, **m**_i_ represents the magnetic dipole moment of particle *i*, *t* the time, γ_o_ the gyromagnetic ratio, and λ the damping constant. An arbitrary vector **b** in the laboratory coordinates can be transferred into the coordinates of particle *i* through:

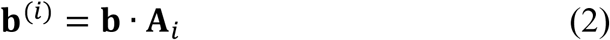

Here, **A**_i_ is the transformation matrix in terms of particle *i*, the computation of which has been discussed in previous studies^44^.

In Equation (1), **B**^(*i*)^_eff_ is contributed by four specified fields. The first one is the external alternating magnetic field, i.e., **B**_ac_ = *B*_o_ cos(2π*ft*) **i**_z_, which is applied along the z-direction of the laboratory coordinates (indicated by **i**_z_) and with the field amplitude *B*_o_ and frequency *f*. The second field is due to the long-ranged magnetic dipole-dipole interaction. It is written in the form of:

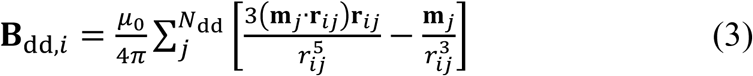

where μ_o_ represents the vacuum permeability, *N*_dd_ the number of neighboring particles of particle *i*, and **r**_ij_ the vector of center-to-center distance (from **m**_j_ to **m**_i_), and *r_ij_* the norm of **r**_ij_. The third field is originated from the magnetocrystalline anisotropy energy barrier, **B**_ani,i_. In present work, we assumed it has a cubic shape that is typical of magnetite. Its expression and comparison with the uniaxial-shaped anisotropy have been detailed previously^44,45^. The last field is due to thermal fluctuation, **B**_therm,i_. By applying the fluctuation-dissipation theorem, **B**_therm,i_ is characterized by zero mean and covariance^46^:

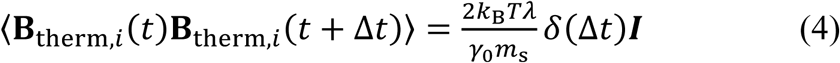

where Δ*t* denotes the discrete time interval, *k*_B_ the Boltzmann constant, *T* the absolute temperature, *m_s_* the magnitude of dipole moment, δ the Dirac delta function implying the white noise field, and **I** the identity matrix.

On the other hand, for spherical MNPs that are freely suspended in viscous liquids. They experience the forces and torques due to Stokes drag (**F**_S,*i*_ and **T**_S,*i*_), magnetic dipole-dipole interaction (**F**_dd,*i*_), osmotic force caused by the steric volume of surfactant polymers (**F**_osm,*i*_), magnetocrystalline anisotropy energy (**T**_ani,*i*_), and thermal agitations (**F**_B,i_ and **T**_B,i_). The neglect of inertia and hydrodynamic interactions are justified for nano-scaled particle size and dominance of the long-ranged magnetic dipole-dipole interactions^47^, respectively. Therefore, the balance equations of force and torque on particle *i* can be written in the form of:

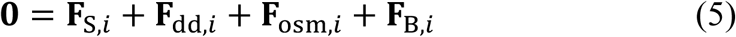

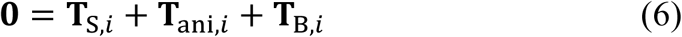

In a quiescent fluid, the Stokes force and torque are given by:

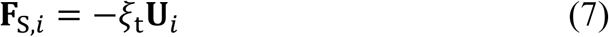

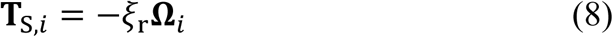

respectively, where the translational and rotational friction coefficients ξ_t_ = 6πη_o_*a* and ξ_r_ = 8πη_o_*a*^3^, respectively, η_o_ represents the fluid viscosity, *a* the particle radius, and **U**_i_ and **Ω**_i_ the velocity and angular velocity of particle *i*. The magnetic dipolar force on particle *i* is given by:

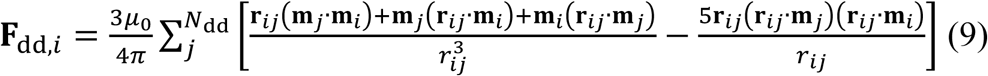

When two particles get close, short-range repulsion, such as the steric force due to the lipid bilayers and electrostatic force, become essential. In current work, we modeled such a repulsion through the osmotic force^48^:

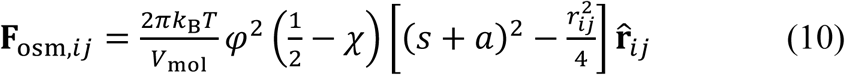

where φ is the volume fraction of polymer in the outer shell, *V*_mol_ the molecular weight of water, χ the Flory-Huggins parameter, and *s* the shell thickness. In Equation (6), **T**_ani,*i*_ is the key term to couple the dynamics of internal magnetic dipole moment and host particle. It indicates the torque due to the deviation of the magnetic dipole moment from the easy axes of particle *i* and reads:

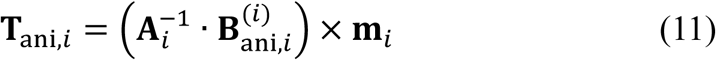

Moreover, because the particles are always colliding with solvent molecules, they are subjected to Brownian force and torque, which are characterized by a Gaussian distribution with zero mean and covariance:

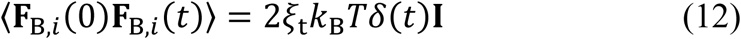

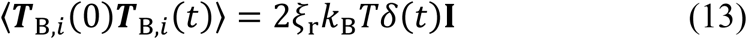

Simulations were carried out for 1000 uniform spherical MNPs with domain magnetization *M*_d_ = 4.46 × 10^S^ A/m and magnetocrystalline anisotropy constant *K*_c_ = 13.5 kJ/m^-3^ (for cubic anisotropy). The temperature and solvent viscosity are fixed at 298.15 *K* and η_o_ = 0.893 mPa · s, respectively. The damping constant was set to λ = 1, although changing the value does not affect the dynamics but require smaller and more time steps^45,49^. For the osmotic force, we set the non-dimensionalized parameter π*a*^3^φ^2^(1/2 − χ)/*V*_mol_ = 5000 in order to produce strong steric repulsion relative to magnetic dipolar attraction. We varied the diameter of magnetic core in the range of 20 nm ≤ 2*a* ≤ 55 nm, shell thickness in the range of 2 nm ≤ *s* ≤ 40 nm, amplitude of the alternating magnetic field in the range of 1 mT ≤ *B*_o_ ≤ 40 mT, and field frequency in the range of 1 kHz ≤ *f* ≤ 40 kHz. By mimicking the experimental scenario, runs were carried out from random particle configurations and under a static magnetic field of intensity 80 mT (along the z-direction). Immediately after such a chaining preprocess, the alternating magnetic fields replaced and sustained for ten periods. The minimum time interval Δ*t* ranges from 1 ps to 1 ns, depending on the particle size yet field frequency. By examining the particle configuration and time evolution of system magnetization, we verified that steady states were typically attained after five field periods and the minimum time interval was delicate enough to capture subtle magnetization dynamics.

### MPS Experiment

MPS measurements were conducted on the 1D-MPI & MPS system with gradient field turned off. 100 µL of MAGiC or MNPs sample with a concentration of 0.1 mg Fe/mL was placed in a 500 µL Eppendorf tube and measured. The excitation field strength and frequency were adjusted in the range of 14-60 mT and 1-35 kHz, respectively. M-H curve and PSF were generated based on the MPS results.

### M-H curve

The excitation magnetic field H and the signal *s*(*t*) over one excitation cycle were extracted, and the magnetization was calculated using

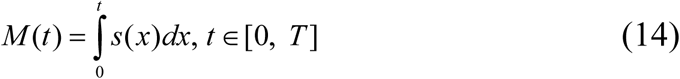

where *M* (*t*) is the magnetization, *T* is the period of the excitation magnetic field. M-H curve was plotted with H as the x-axis and M as the y-axis.

### PSF

The MPS signal was processed using Equation (S2), where the excitation field *H*(*t*) was used as *x_s_*(*t*). The resulting *IMG* was plotted against *H*(*t*) to generate the PSF. The physical distance *_x_* can be calculated using the gradient value G:

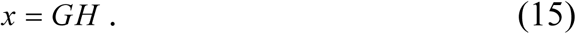

PSF of different tracers were normalized to allow for comparison. FWHM was calculated from the PSF using:

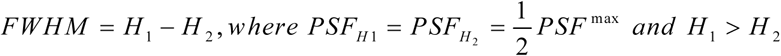

*PSF_H_* represents the value of PSF when the external magnetic field is *H*, and *PSF*^max^ is the maximum value of PSF.

### MPI imaging of phantoms

1D phantoms were designed, 3D-printed, and filled with 0.2 mg Fe/mL MAGiC or other MNPs for imaging. Images were acquired with the 1D-MPI & MPS system with an excitation field of 1kHz and 40 mT.

2D and 3D phantoms were designed, 3D-printed, and filled with 0.2 mg Fe/mL MAGiC or other MNPs for imaging. Phantoms filled with MAGiC were imaged using the MAGiC-dedicated system. Since the excitation parameters of the MAGiC dedicated system were not ideal for VivoTrax+™ and synomag®-D, phantoms filled with these MNPs were imaged with the Momentum scanner (Momentum, Magnetic Insight, Alameda, USA) to allow for fair comparison. Default scan mode was used, with a gradient of 5.7 T/m in both the x and y directions.

### Animals

Rats and mice were purchased from Beijing Vital River Laboratory Animal Technology Co. and maintained in an animal facility at Institute of Automation, Chinese Academy of Sciences. All animal experiments were performed in accordance with the Declaration of Helsinki and all procedures were approved by the ethics committee of Institute of Automation (IA21-2311-010201).

### In vivo imaging of MAGiC

To demonstrate in vivo imaging capability of MAGiC in deep tissue, 100 g female Sprague Dawley rats were used for imaging. Prior to imaging, rats were fasted for 12 hours. 10 µL of 5 mg Fe/mL MAGiC was encapsulated in an enteric capsule. The capsule was administered via oral gavage. After oral administration, rats were imaged with the MAGiC dedicated MPI system at 0 h, 1 h, 3 h, 5 h, 7 h, 9 h, and 24 h. Anatomical images were acquired with a custom-built CT scanner and were registered with the MPI images.

### In magnetic actuation of MAGiC

To study the magnetic enrichment and actuation behavior of MAGiC, phantoms were filled with 0.02 mg Fe/mL MAGiC solution. MAGiCs were magnetically actuated or enriched using a 5 mm × 5 mm × 10 mm N52 magnet. To study the upstream locomotion of MAGiC, a peristaltic pump (BT100-2J, LongPump, China) was used to propel the fluid at a flow rate of 20 mL/min through a quartz glass tube with an inner diameter of 4 mm. MAGiC was magnetically actuated using a cylindrical N52 magnet with a diameter of 3 mm and a height of 2.1 mm.

### In vivo tracking of MAGiC locomotion

To track MAGiC locomotion with MPI in vivo, 100 g female Sprague Dawley rats were intraperitoneally injected with 100 µL MAGiC (0.5 mg Fe/mL). To examine the initial distribution of the particles, MPI images were acquired 5 minutes after administration. Subsequently, two rounds of magnetic navigation were performed. For the first round, a 5 mm × 5 mm × 10 mm N52 magnet was placed at target position #1 to collect the diffusely distributed MAGiCs. MPI images were acquired at 80 minutes to examine the accumulation of MAGiCs. For the second round, the same magnet was repositioned to target position #2 to induce locomotion of MAGiCs. MPI images were acquired at 80 min and 220 min to track the locomotion. CT images were acquired with the custom-built scanner and registered with MPI images.

### Biocompatibility studies

To examine the biocompatibility of MAGiC, 6-week-old female ICR mice were randomly assigned to experimental or control groups. Mice in the experimental groups were administered 100 µL 0.5 mg Fe/mL MAGiCs either via oral gavage or via intraperitoneal injection. Mice in the control groups were administered 100 µL PBS. To test the liver and renal functions, blood samples were collected from the orbital venous plexus at either 24 h or 7 days after administration. Serum was prepared and liver function parameters ALT, AST, TBIL, ALB, and renal function parameters BUN/UREA and CR were analyzed with an automatic biochemical analyzer. To perform the histological analysis, mice were euthanized at 24 h or 7 days after administration, major organs and different parts of the GI tract were extracted and examined with H&E.

## Notes

### Competing Interest Statement

The authors have declared no competing interest.

