## supplementary material for "Magnetically induced magnetosome chain (MAGiC): A biogenic magnetic-particle-imaging tracer with high performance and navigability"

**Comparison with existing simulation models**

Park et al. have numerically predicted the magnetization reversals of magnetosome arrays, based on the delicate model of ten octahedron MNPs that are immobilized in straight chains[1]. The outer lipid bilayer, as a non-negligible component of magnetosomes, was taken into account by setting a consistent surface separation of 5 nm between the nearest neighbors. However, this is not the case for the magnetosomes that are freely suspended in solvent such as water, which may assemble into chains with various lengths and with larger inter-particle separations due to the swelling effect of the polymer shell. Such a fact was examined in our experimental characterizations. The magnetosomes have a crystal around 27 nm and a membrane around 3.6 nm. However, the particle surface separation becomes around 30 nm.

In order to account for the precession of internal magnetic dipole moment and simultaneously the physical rotation of MNP, prior works have incorporated the Landau-Lifshitz-Gilbert (LLG) equation into Fokker-Planck equation[2,3] or Langevin dynamics simulations[4–6]. However, the focus of such work is only on the rotational rather than translational degrees of freedom of the particles, leading to the absent investigation of chaining dynamics.

Therefore, existing simulation models cannot meet the simulation requirements for MAGiC. We developed algorithms that couple the LLG equation into Brownian dynamics simulations, accounting for both rotational and translational motion of the magnetosomes. For details, please refer to the Methods section.

**Preparation of magnetosome chains connected by cytoskeletal filaments**

After fully resuspending the collected bacterial cells in 10 mM PBS in a beaker, they were disrupted using an ultrasonic cell crusher, specifically with the following settings: 300 W power output, 3 s working time, 5 s working interval, and a total working time of 30 min. After ultrasonication, the beaker was placed on a magnet at 4 °C overnight for adsorption. The supernatant was discarded, and the adsorbed sediments were resuspended in PBS. They were then cleaned using ultrasonication at 80 W power for 20 min and collected again using the magnet. The cleaning steps were repeated, with the ultrasonic power decreasing with each iteration. This resuspension-ultrasonication process was repeated 7–8 times until the protein concentration of the supernatant was lower than 0.1 mg/mL, at which point magnetosome purification was considered complete.

**MAGiC dedicated 3D MPI scanner**

The dedicated MAGiC MPI system adopted a field-free point (FFP) based design. The FFP was generated using a pair of NdFeB permanent magnets, with gradient strengths of 1.25 T/m in the x and z directions and

2.5 T/m in the y direction. A solenoid with a winding diameter of 45 mm and a length of 100 mm was used to generate the excitation magnetic field in the x-direction, which was wound with 180 turns of 0.1 mm × 350 litz wire. An external excitation coil (for background signal compensation) with the same size and winding was connected in series with the excitation solenoid. Additionally, a 14 μF capacitor group was connected in series to achieve resonance at 1 kHz. The excitation circuit was driven by an AE Techron 7224 (AE Techron Elkhart, USA) amplifier, generating an excitation magnetic field of 1 kHz and 40 mT. A pair of Helmholtz coils generated the focus field in the y-direction. Each coil had a winding diameter of 154 mm, a thickness of 42 mm, and was wound with 350 turns of 2 mm enameled wire. The distance between the centers of the two coils was 110 mm. The focus coils were driven by another AE Techron 7224 amplifier, generating a focusing field of 10 Hz and 45 mT. Another pair of Helmholtz coils generated the focus field in the z-direction. Each coil had a winding diameter of 292 mm, a thickness of 38 mm, and was wound with 280 turns of 2 mm enameled wire. The distance between the centers of the two coils was 177 mm. The focus coils were driven by an AE Techron 7548 (AE Techron Elkhart, USA) amplifier, generating a focusing field of 0.5 Hz and 25 mT. The receive coil was an x-direction solenoid with 39 mm in diameter and 29 mm in length, wound with 114 turns of 0.2 mm enameled wire. An compensation coil with the same structure was placed inside the external excitation coil and was differentially connected with the receive coil. The signal received from the coils was passed through a 2.2 kHz high-pass filter (EF 113, Thorlabs, New Jersey, USA) and a 100 kHz low-pass filter (EF 502, Thorlabs, New Jersey, USA), and was amplified by a low-noise pre-amplifier (SR560, SRS, Sunnyvale, CA, USA) before being converted to digital signal by an ADC (PCIe-6374, National Instruments, USA). The control signals for both the excitation and focus fields were generated by PCIe-6374 (using two in total). This system provided a FOV of 6.5 * 3.5 * 3.5 $cm^3$. 2D imaging speed was 10 frames per second and 3D imaging speed was 0.5 volume per second.

**1D MPI & MPS scanner**

1D-MPI & MPS system employed a 4 T/m gradient field in the x direction, which was generated by a pair of Helmholtz coils driven by an AE Techron 7548 amplifier. The gradient coils were 70 mm in diameter and spaced 80 mm apart, each wound with 700 turns of 2.24 mm enamel-coated wire. The gradient field can be turned off for MPS measurement. The excitation field was generated by a solenoid with 26 mm in diameter and 100 mm in length, wound with 160 turns of 0.1 mm × 300 litz wire. The excitation coils were driven by an AE Techron 7224 amplifier, generating excitation fields with frequency in the range of 0-50 kHz and strength in the range of 0-60 mT. A solenoid gradiometer receive coil was used. The receive and compensation sections were positioned symmetrically around the center with 24 mm apart. Each section was 19 mm in diameter and 12 mm in length, wound with 232 turns of 0.2 mm enameled wire. The received signal was passed through a 240 kHz low-pass filter (EF 504) and amplified by a low-noise pre-amplifier (SR560) before being converted to a digital signal by an ADC (USB 6356, National Instruments, USA). The control signals for both the excitation and gradient fields were generated by USB 6356. This system provided a 1D FOV of 2 cm when used as 1D MPI. The MPS scanning and 1D imaging speed was 1000 frames per second (when the excitation frequency is 1kHz). The minimum detection voltage of $10^{-4}$ V when this system is used for MPS.

**MAGiC image reconstruction**

The x-space reconstruction formula is[7]:

$$IMG(x_s(t)) = \frac{s(t)}{B_1 mkG\dot{x}_s(t)},\ k = \frac{\mu_0 m}{k_B T^p}, \tag{S1}$$

where $IMG$ is the reconstructed image, $x_s(t)$ is the position of the FFP, $s(t)$ is the time domain signal, $\dot{x}_s(t)$ is the speed of FFP, $B_1$ is the sensitivity of the receive coil, $m$ is the magnetic moment of the particle, $G$ is the field gradient, $\mu_0$ is the permeability of free space, $k_B$ is the Boltzmann constant, $T^p$ denotes the particle temperature.

Due to the coercivity of MAGiC (Fig. 1(b)), Equation (1) was modified and a half-cycle x-space algorithm was proposed:

$$IMG(x_s(t_{\dot{x}_s>0}) - \frac{H_C}{G}) = \frac{s(t_{\dot{x}_s>0})}{B_1 mkG\dot{x}_s(t_{\dot{x}_s>0})} \tag{S2}$$

where $H_C$ represents the coercivity of MAGiC, $\dot{x}_s > 0$ represents that only the signal received during the forward movement of the FFP is used for reconstruction, $-\frac{H_C}{G}$ is used to correct the positional deviation caused by the coercivity.

### Image quantification

Regions of interests (ROI) were selected and pixel values of pixels within the ROI were summed and normalized. Finally, the quantification value is calculated based on the linear relationship between the summed value and the iron content.

### Magnetosome membrane removal

To remove the magnetosome membrane, magnetosomes were collected by magnetic collection, re-suspended in 2 M NaOH with 10% SDS and boiled for 10-15 min. Magnetosomes without membranes were then washed with PBS and collected with a magnet.

### Magnetosome membrane protein removal

To remove the magnetosome membrane proteins, magnetosomes were collected by magnetic collection, resuspended in 2 mg/mL proteinase K (Biotopped, Los Angeles, America, CAS#39450-01-6) and incubated at 56 °C for 3 h. Magnetosomes without membrane proteins were then washed with PBS and collected with a magnet.

**sodium lactate medium**

| Compound | Amount（1L） |
|---|---|
| sodium lactate（55%-65%） | 2mL |
| $NH_4Cl$ | 0.4g |
| $K_2HPO_4$ | 0.5g |
| $MgSO_4 \cdot 7H_2O$ | 0.1g |
| Yeast | 0.1g |
| Sodium thioglycolate | 0.05g |
| Mineral solution（10×） | 0.5mL |

Dissolve ingredients in the order given and adjust pH to 7.0 with HCl. Steam sterilized at 121°C.

**Mineral solution（10×）**

| Compound | Amount（1L） |
|---|---|
| Nitrilotriacetic acid | 15g |
| MgSO4·7H2O | 30g |
| MnSO4·2H2O | 5g |
| NaCl | 10g |
| FeSO4·7H2O | 1g |
| CoSO4·7H2O | 1.8g |
| CaCl2·2H2O | 1g |
| ZnSO4·7H2O | 1.8g |
| CuSO4·5H2O | 0.1g |
| [KAl(SO4)2·12H2O] | 0.2g |
| H3BO3 | 0.1g |
| Na2MoO4·2H2O | 0.1g |
| NiCl2·6H2O | 0.25g |
| Na2SeO3·5H2O | 0.003g |

First dissolve nitrilotriacetic acid with NaOH，then add minerals，Steam sterilized at 121°C，Store at 4°C.

**Fermentation medium**

| Compound | Amount（4.5L） |
|---|---|
| sodium lactate（55%-65%） | 4.0g |
| $NH_4Cl$ | 1.0g |
| $K_2HPO_4 \cdot 3H_2O$ | 3.0g<br>（Autoclavable individually） |
| $MgSO_4 \cdot 7H_2O$ | 1.2g |
| Yeast | 3.0g |
| Mineral solution（10×） | 3.5mL |

**Feed medium**

| Compound | Amount（0.5L） |
| --- | --- |
| sodium lactate（55%-65%） | 100g |
| $NH_3 \cdot H_2O$ | 18mL |
| $MgSO_4 \cdot 7H_2O$ | 1.2g |
| Yeast | 3.0g |
| Mineral solution（10×） | 3.5mL |
| $FeCl_3 \cdot 6H_2O$ | 2.0g |

### Supplementary figures & table

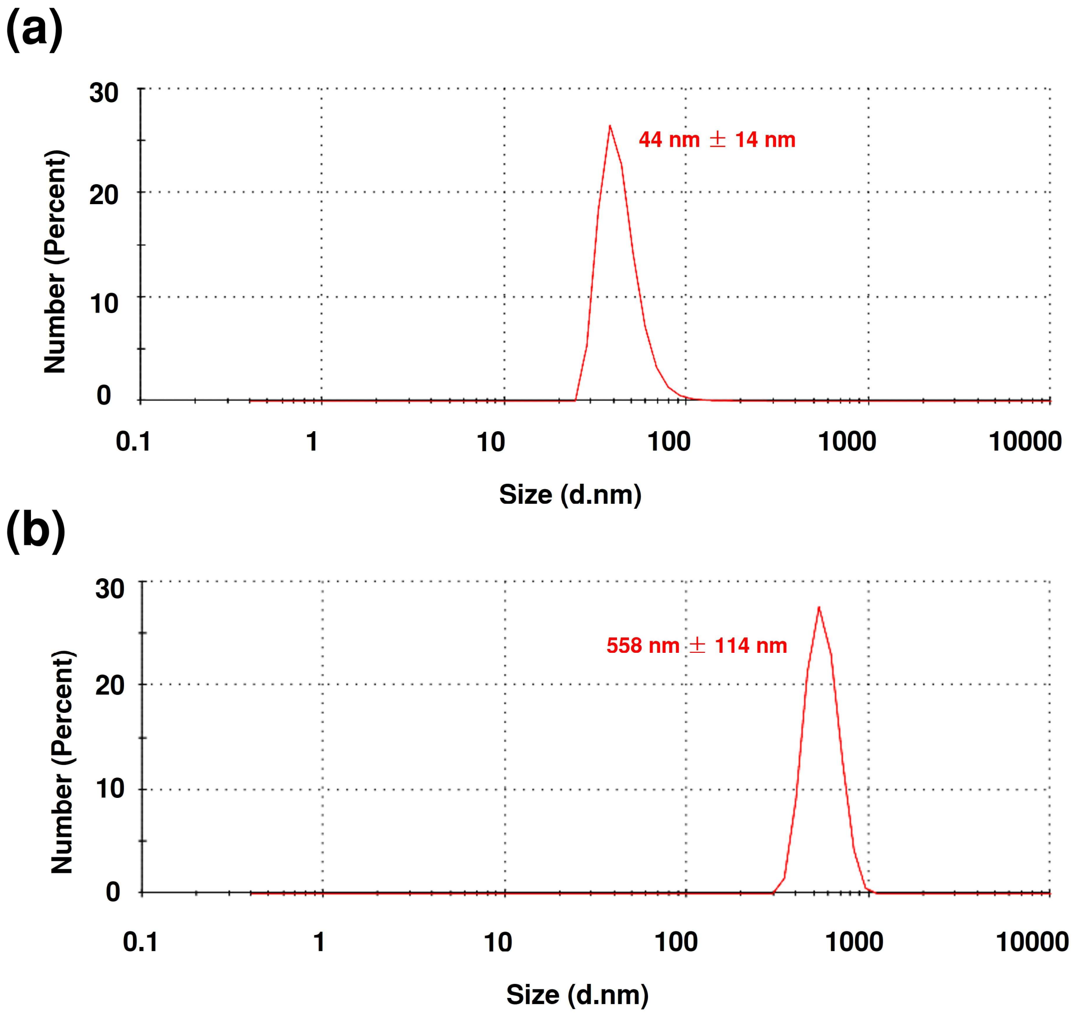


Fig. S1. The hydrodynamic diameter of dispersed magnetosomes (a) and MAGiCs (b). Based on the hydrodynamic diameter values, it is estimated that each MAGiC contains approximately 10-15 magnetosomes.

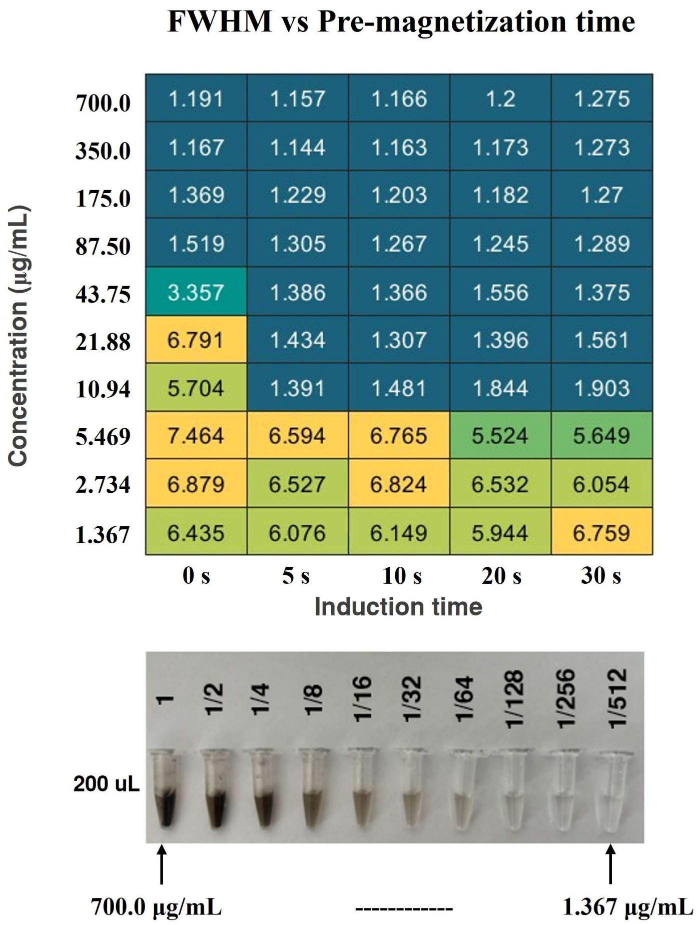


Fig. S2. The effect of magnetic induction time on MAGiC's PSF FWHM. Upon induction with an external magnetic field, magnetosomes transform into MAGiCs. The chart shows the FWHM of MAGiC under different concentrations and induction times. At higher concentrations (87.5 µg/mL and above), there is no significant change in FWHM before and after induction. This indicates that at higher concentrations, magnetosomes can form MAGiCs under the excitation magnetic field, without the need for induction. At concentrations between 43.75 µg/mL and 10.94 µg/mL, the FWHM significantly decreases after induction; however, induction times beyond 5 seconds do not show noticeable differences. This aligns with the simulation results, where magnetosomes can assemble into MAGiC within 500 µs under the induction field. A duration of 5 seconds is sufficient for this process. At concentrations below 10.94 µg/mL, the PSF FWHM significantly higher than those at higher concentrations after induction. This indicates that at low concentrations, the chain formation process of magnetosomes is restricted, which affects the performance of MAGiC.

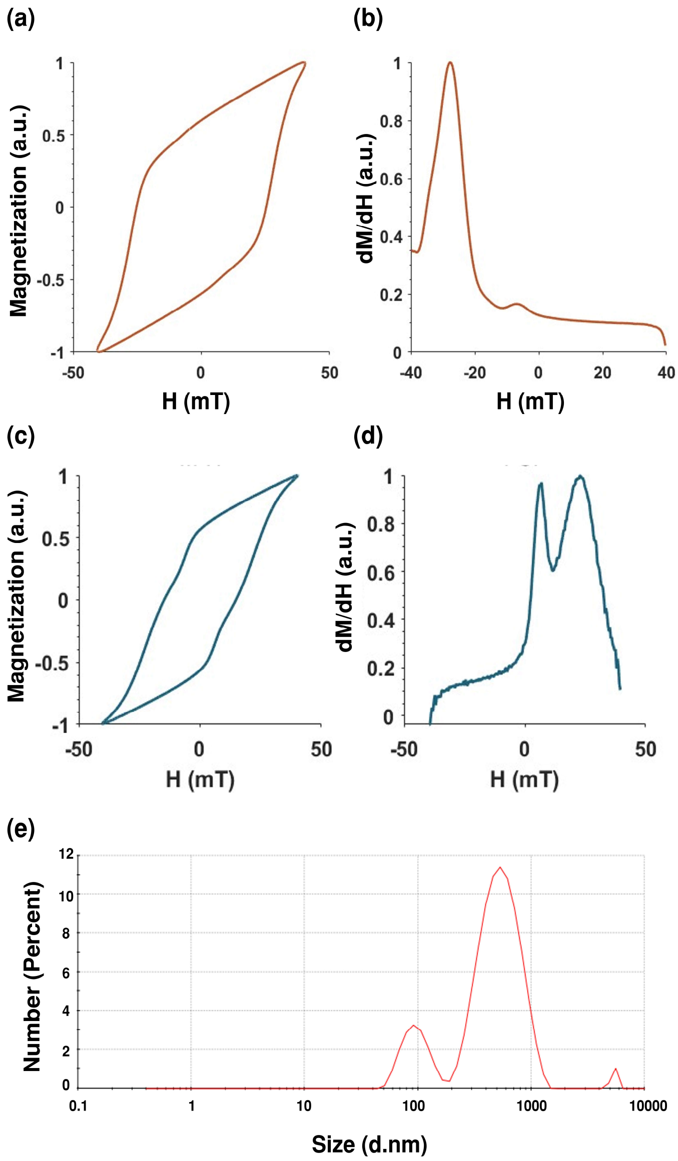


Fig S3. The dynamic magnetization curve and PSF of ZC3M25 bacteria and extracted magnetosome chains linked by cytoskeletal filaments. (a) The dynamic magnetization curve and (b) PSF of ZC3M25 bacteria. (c) The dynamic magnetization curve and (d) PSF of extracted magnetosome chains connected by cytoskeletal filaments; the PSF shows two peaks, both of which are relatively broad compared to MAGiC. (e) The hydrodynamic size distribution of extracted magnetosomes chains; the three distribution peaks correspond to 98.6 nm (16.1%), 562.1 nm (82.6%), and 5405 nm (1.3%). This indicates that we successfully extracted magnetosome chains with a primary hydrodynamic size of approximately 562 nm (containing about 15-25 magnetosomes) and some shorter and longer chains. The two peaks observed in the PSF mainly originate from the 562.1 nm and 98.6 nm magnetosome.

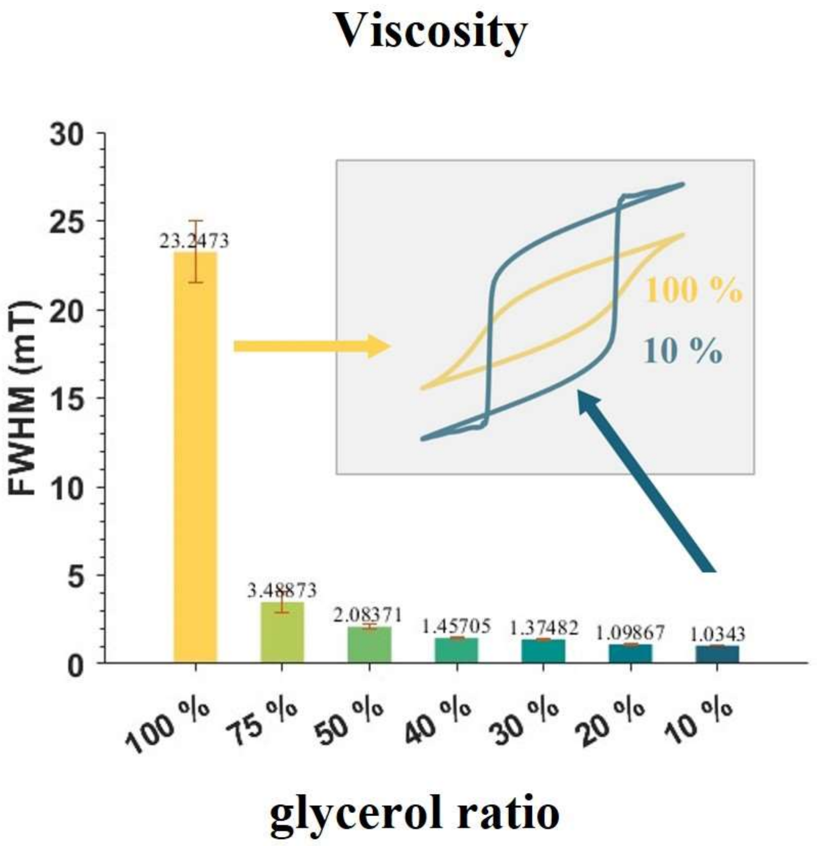


Fig. S4. PSF FWHM of MAGiC in viscous medium. As the volume ratio of glycerol gradually increases, the FWHM of MAGiC's PSF also increases, indicating a deterioration in imaging quality.

Table S1. MPS performance of MAGiC in comparison with commercially available MPI tracers under sinusoidal excitation at a frequency of 1kHz and an amplitude of 40 mT.

| | **Resolution (1 T/m gradient field)** | **Signal amplitude(a.u.)** |
|---|---|---|
| MAGiC | 0.6 mm | 91.0 |
| VivoTrax+™ | 15.3 mm | 1.0 |
| synomag®-D | 11.0 mm | 3.2 |
| Perimag® | 7.6 mm | 1.3 |

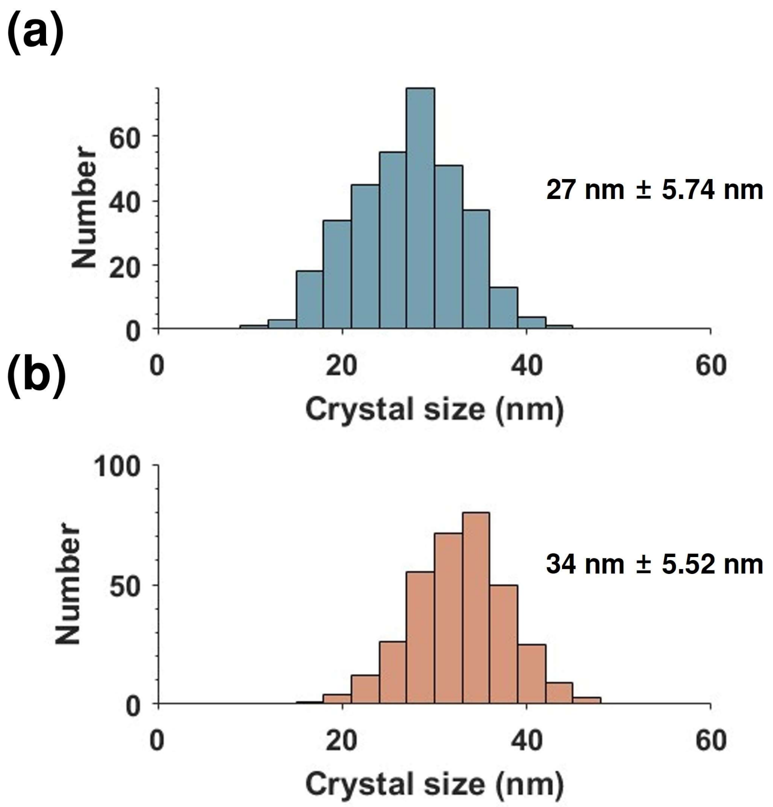


Fig. S5. The crystal size of magnetosomes measured in TEM. A total of 330 magnetosome size data were collected from five TEM images (each group). (a) The magnetosomes extracted from ZC3M25 cultured in screw-capped bottles, showing a mean crystal size of 27 nm. (b) The magnetosomes extracted from ZC3M25 cultured in fermenters, demonstrating a mean crystal size of 34 nm.

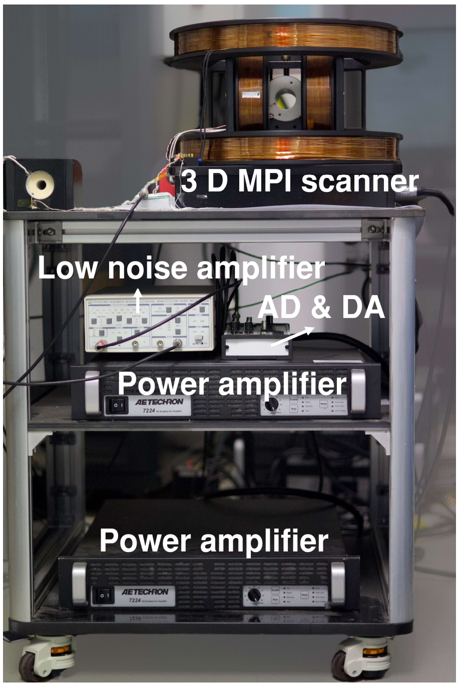


Fig. S6. MAGiC dedicated 3D MPI scanner

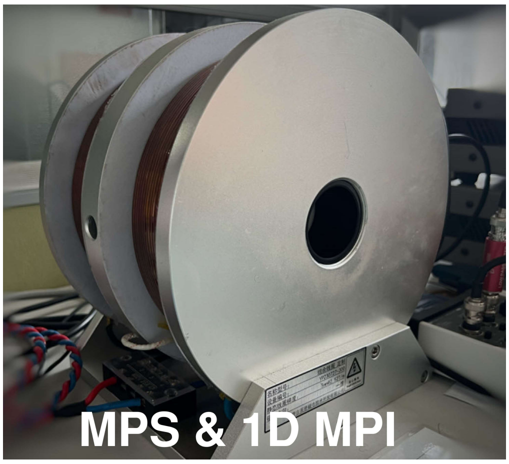


Fig. S7. 1D MPI & MPS scanner for MAGiC

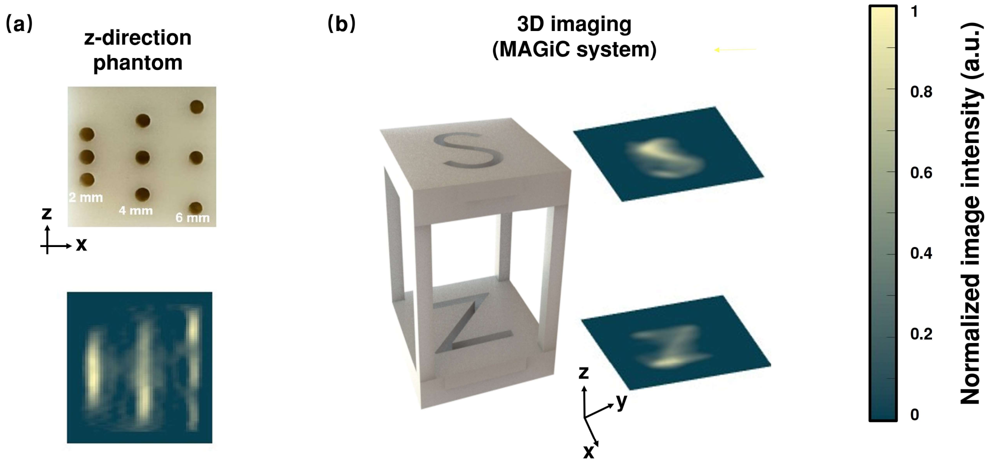


Fig. S8. MAGiC's z-axis resolution and 3D imaging result using MAGiC dedicated MPI. (a) The imaging results of the z-axis resolution phantom show that MAGiC achieves 6 mm resolution in the z-axis. (b) The 3D imaging results of MAGiC.

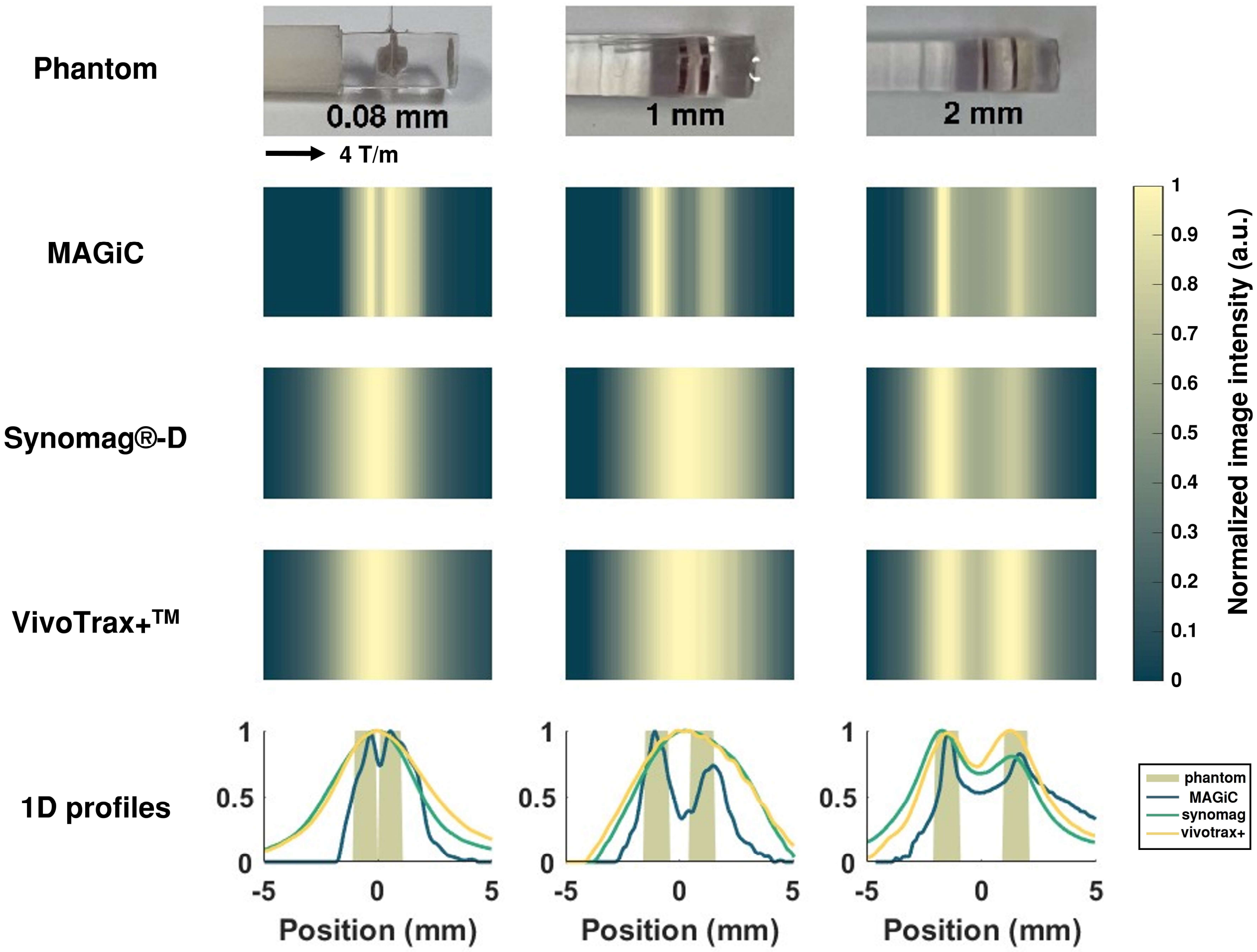


Fig. S9. The imaging results of resolution phantom in 4 T/m 1D MPI. The spatial resolution of MAGiC reached 0.08 mm, while those of synomag®-D and VivoTrax+$^{TM}$ were around 2 mm. The 0.08 mm resolution phantom and MAGiC imaging result are discussed in details in Fig. S10.

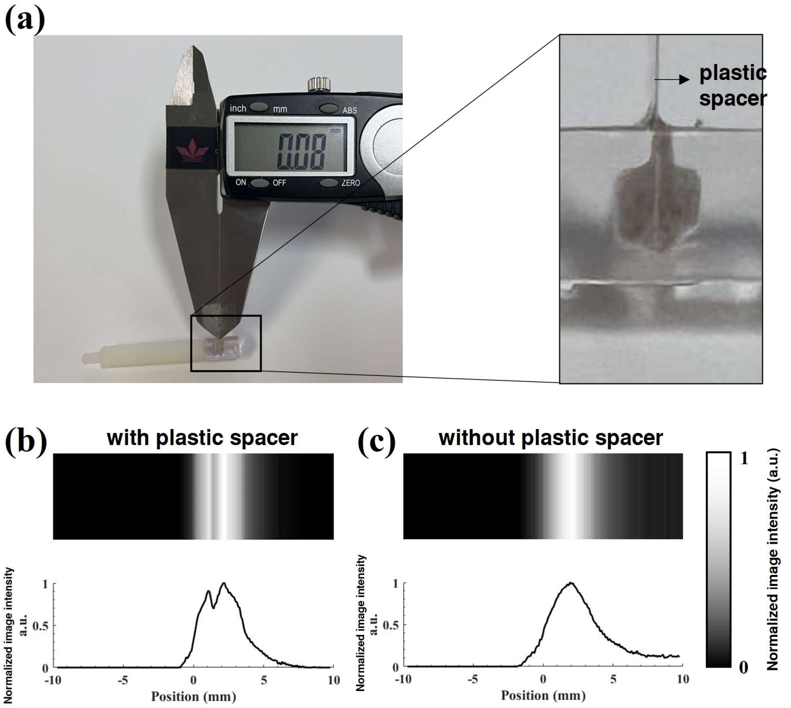


Fig. S10. Photograph of the 0.08 mm resolution phantom and MAGiC imaging result. (a) The 0.08 mm resolution phantom was constructed by inserting a 0.08 mm thick plastic spacer into a sample chamber. We conducted imaging with (b) and without (c) the plastic spacer. The double-peak spectrum observed in (b) and the single-peak in (c) confirm that MAGiC provides a spatial resolution of 0.08 mm.

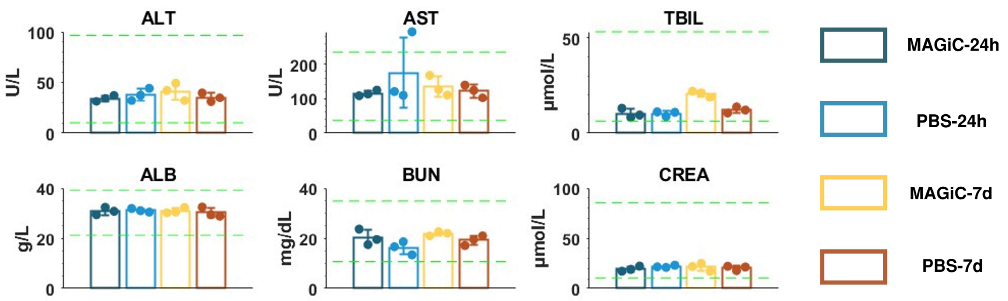


Fig. S11. Liver and renal function tests after oral gavage of MAGiC. Green dashed lines indicate the normal range.

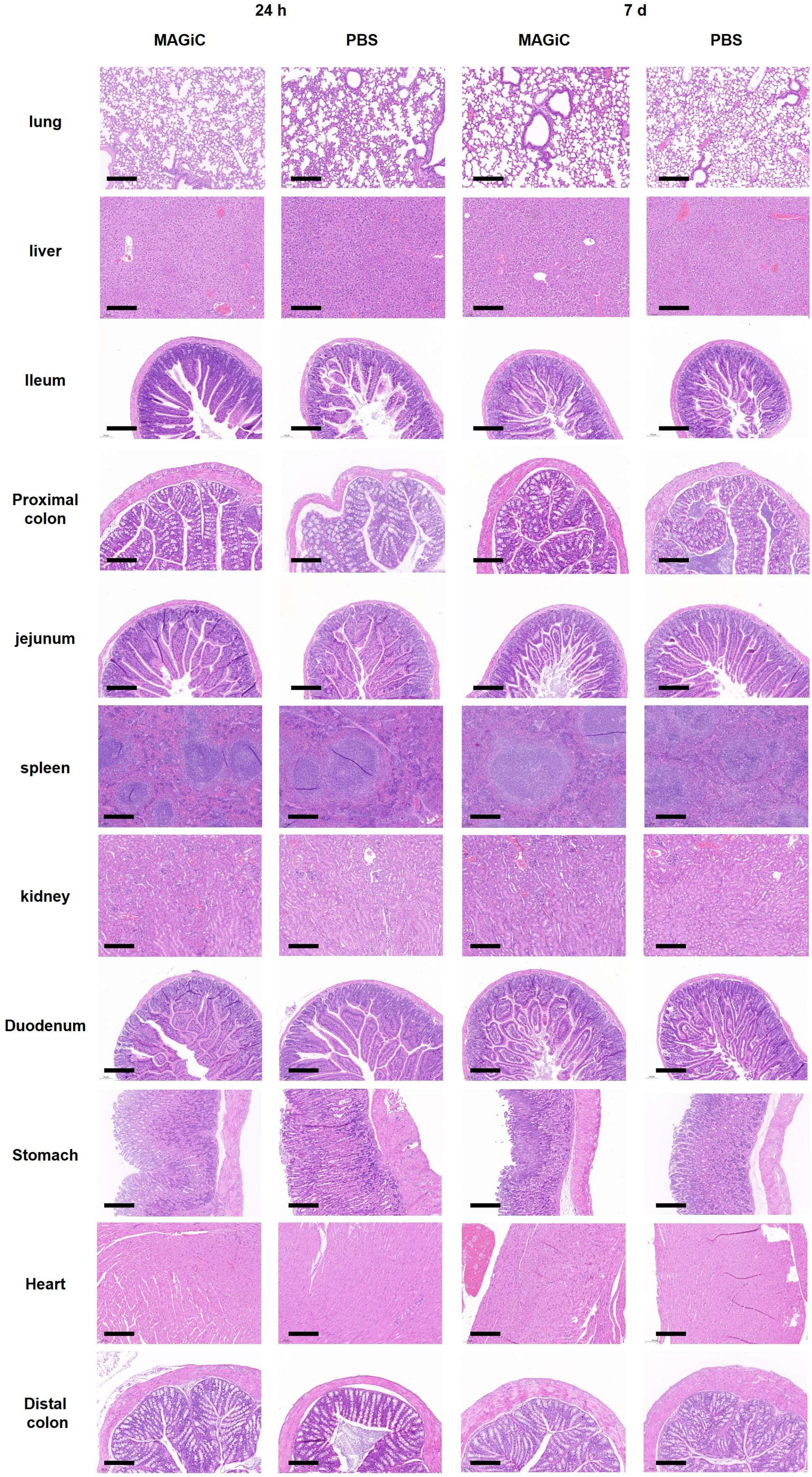


Fig. S12. Histopathological sections of major organs after oral gavage of MAGiC. Scale bar: 200 μm

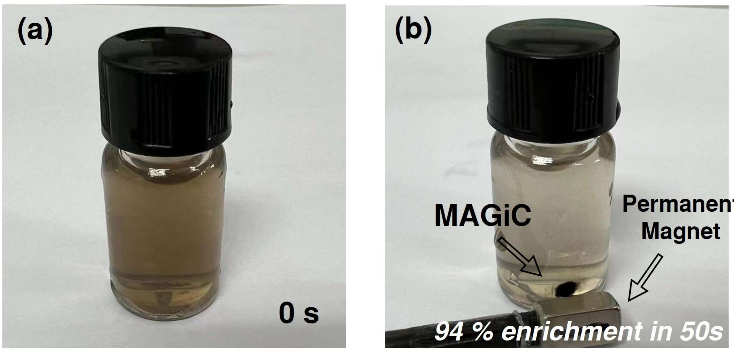


Fig. S13. The enrichment of MAGiC under the field created by a permanent magnet.

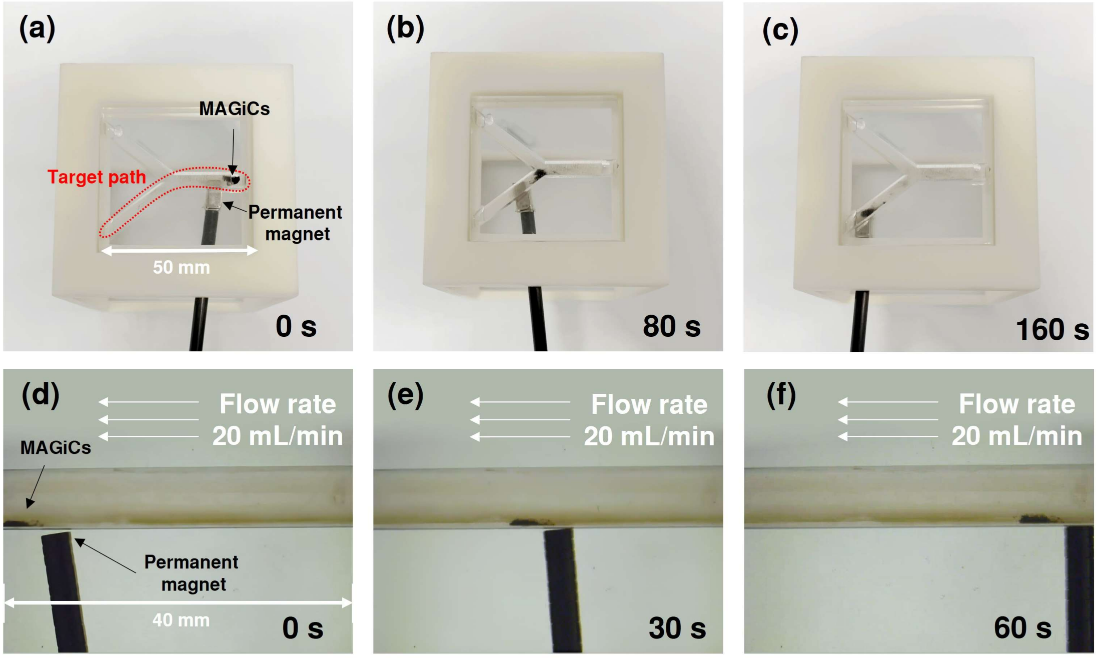


Fig. S14. In vitro magnetic navigation of MAGiC. (a)-(c) Locomotion of a cluster of MAGiC along the predetermined path in a phantom. (d)-(f) Upstream navigation of MAGiC under a flow rate of 20 mL/min.

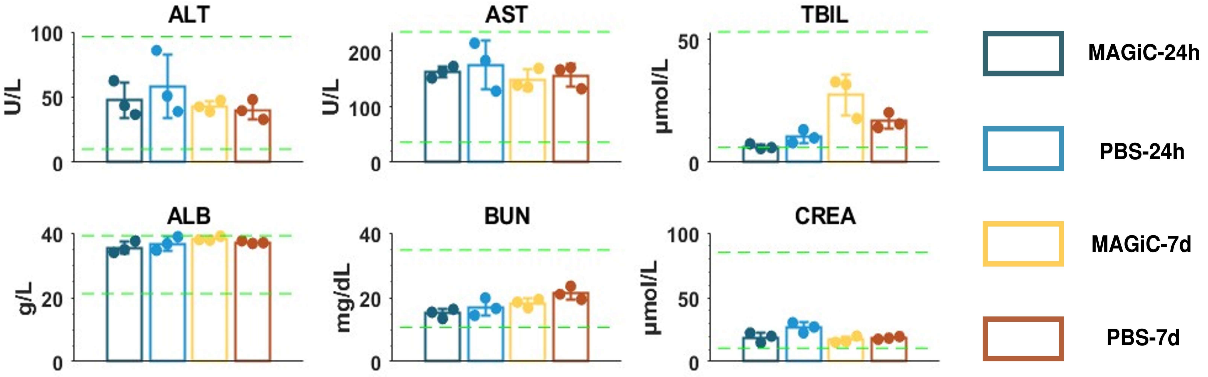


Fig. S15. Liver and renal function tests after intraperitoneal injection of MAGiCs.

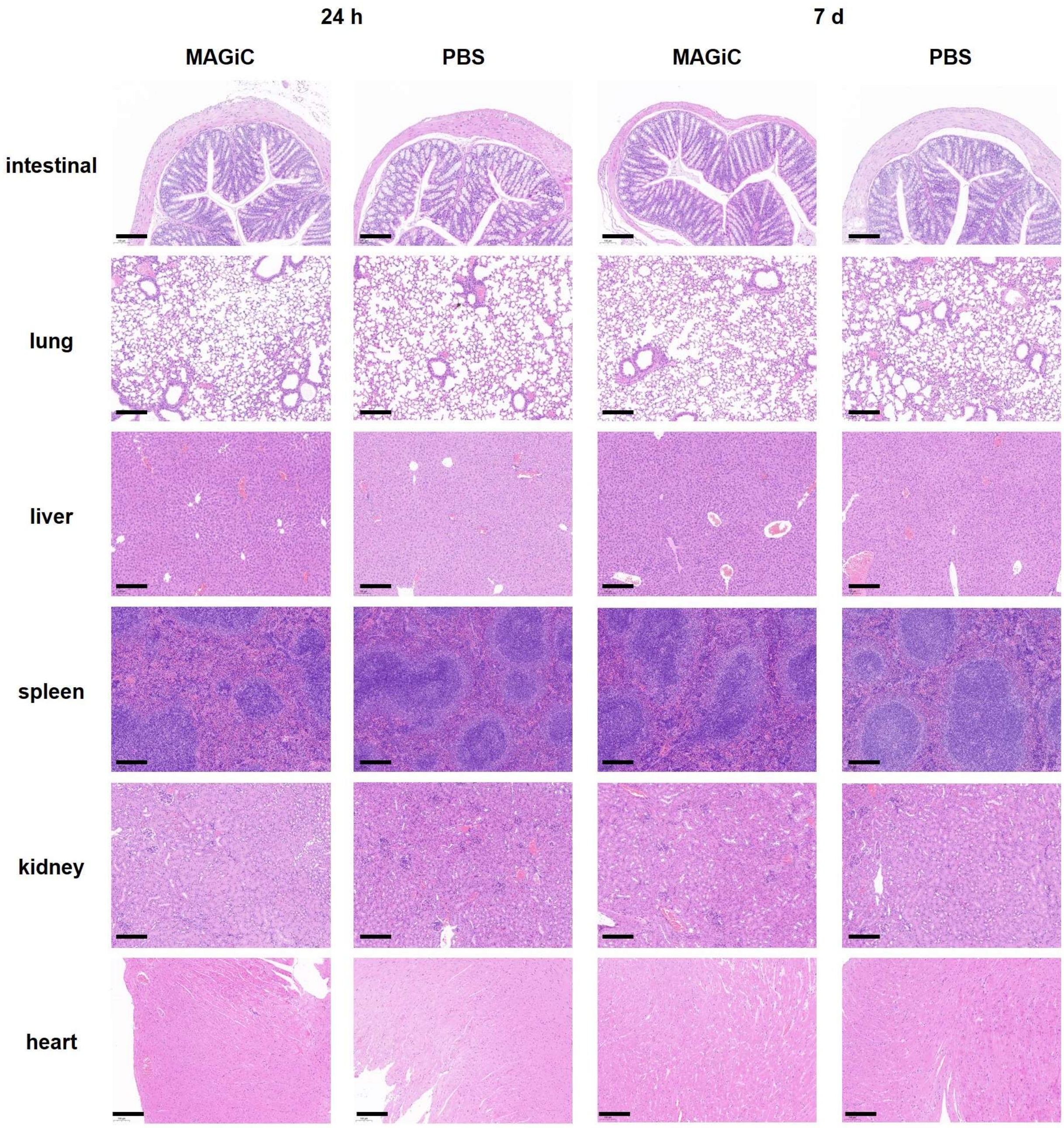


Fig. S16. Histopathological sections of major organs after intraperitoneal injection of MAGiCs. Scale bar: 200 μm

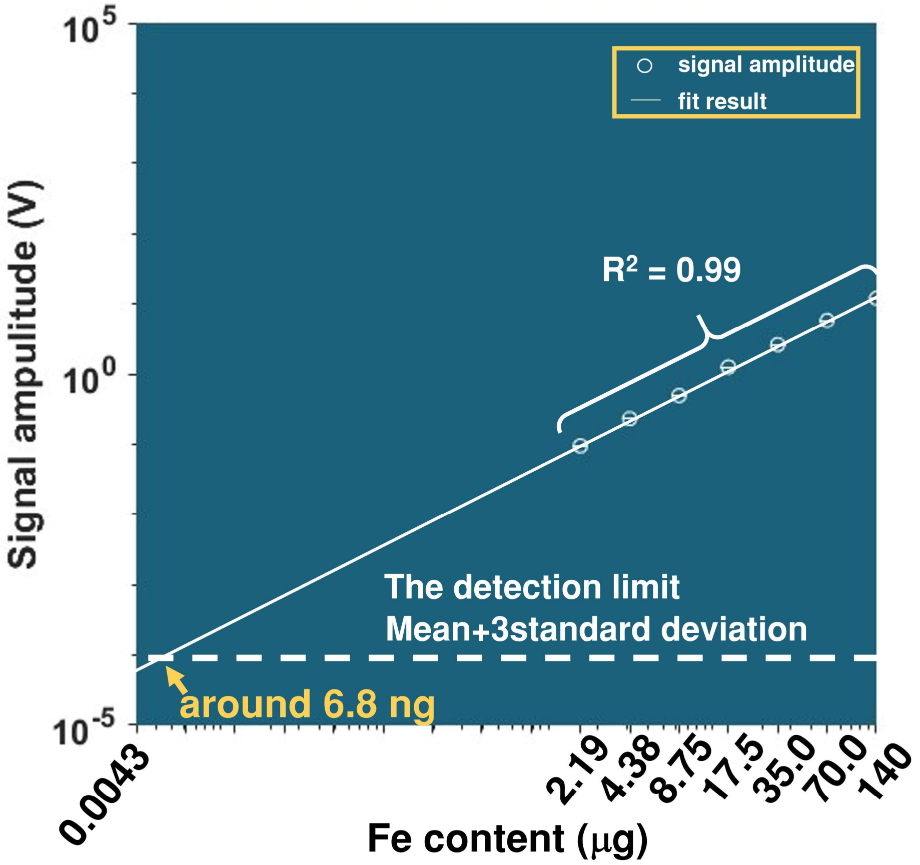


Fig. S17. The relationship between signal intensity and the content of MAGiC. The signal intensity of MAGiC shows a linear relationship with its content. Based on the minimum detection limit of $10^{-4}$ V in MPS, theoretically, the minimum detectable amount of MAGiC is 6.8 ng.
